# Multi-omics identification of extracellular components of the fetal monkey and human neocortex

**DOI:** 10.1101/2024.09.25.615077

**Authors:** Felipe Vilicich, Dhanya Vettiatil, Seth Kattapong-Graber, Nadiya Nawsheen, Neel Patel, Alexandra Quezada, Elizabeth Gurney, Emma Smith, Hallie Nelson, Susan Pesci, Jessica Atrio, Nadjeda Moreno, Aragorn Jones, Melinda Murphy, Nerys Benfield, Jon Hennebold, Nita Solanky, Steven Lisgo, Ian Glass, Birth Defects Research Laboratory (BDRL), Simone Sidoli

## Abstract

During development, precursor cells are continuously and intimately interacting with their extracellular environment, which guides their ability to generate functional tissues and organs. Much is known about the development of the neocortex in mammals. This information has largely been derived from histological analyses, heterochronic cell transplants, and genetic manipulations in mice, and to a lesser extent from transcriptomic and histological analyses in humans. However, these approaches have not led to a characterization of the extracellular composition of the developing neocortex in any species. Here, using a combination of single-cell transcriptomic analyses from published datasets, and our proteomics and immunohistofluorescence analyses, we provide a more comprehensive and unbiased picture of the early developing fetal neocortex in humans and non-human primates. Our findings provide a starting point for further hypothesis-driven studies on structural and signaling components in the developing cortex that had previously not been identified.

## INTRODUCTION

During development, precursor cells are constantly engaging with their surrounding extracellular environment, which plays a critical role in guiding their differentiation and functional maturation. These interactions are mediated through various signaling pathways, mechanical forces, and the composition of the extracellular matrix, all of which provide essential cues that direct precursor cells toward specific lineages, provide structural support, and guide cytoarchitecture. The dynamic nature of these interactions ensures that precursor cells adapt to the changing needs of the developing tissue, ultimately contributing to the formation of a fully functional organ. Without the precise and continuous communication between precursor cells and their extracellular environment, the complex process of tissue and organ development would be severely compromised.

Much of our understanding of development, particularly for the mammalian neocortex, comes from extensive research involving several methodologies. Histological analyses, heterochronic and heterotopic cell transplants, and genetic manipulations in non-primate mammals have provided profound insights into the cellular and molecular mechanisms that drive neocortical development (e.g. McConnell, 1995; Leone et al., 2008; Grove and Fukuchi-Shimogori T, 2003; Creig et al., 2013; O’Leary et al., 2007; DeFelipe et al., 2013). These studies have been instrumental in revealing how precursor cells in the neocortex differentiate into various neuronal subtypes and establish the complex architecture of this brain region. Additionally, transcriptomic and histological analyses in humans, although less extensive, have complemented these findings by offering a glimpse into the unique aspects of human neocortical development (e.g. Kostovic, 2020; Braun et al., 2023; Eze et al., 2021; Wang et al., 2024; Zeng et al.,2023; Polioudakis et al., 2019). This multi-faceted approach has been crucial in unraveling the intricacies of neocortical development, highlighting both conserved mechanisms across species and those that may be unique to humans.

While significant progress has been made in understanding neocortical development through histological, genetic, and transcriptomic studies, these approaches have certain limitations. Notably, they have not provided an unbiased or detailed characterization of the extracellular composition within the developing mammalian neocortex. Without a thorough understanding of the extracellular niche and its dynamic interactions with developing cells, our knowledge of how the neocortex forms and functions remains incomplete. This gap in the research highlights the need for more advanced and integrative approaches to fully characterize the extracellular components and their roles in shaping the developing neocortex.

To begin addressing the gaps in our understanding of the extracellular environment in neocortical development, we have employed a multifaceted approach combining single-cell transcriptomic analyses from published datasets with our own proteomics and immunohistofluorescence analyses. This integrated high-resolution methodology has allowed us to generate a more detailed picture of the early developing fetal neocortex in both humans and non-human primates (NHPs). Our findings also reveal previously unidentified structural and signaling components, offering new avenues for hypothesis-driven research. These discoveries lay the groundwork for further exploration into the specific roles these extracellular elements play in guiding neocortical development, potentially leading to a deeper understanding of both normal brain development and neurodevelopmental disorders.

## RESULTS

### Database of human genes encoding extracellular proteins

Several published databases have provided information about extracellular proteins encoded in the human genome. These include Matrisome (Naba et al., 2012), SignalP (Teufel et al., 2022), MDSEC, Spoctopus (Viklund et al., 2008), and Phobius (Kall et al., 2004). These databases were obtained by validation of candidates in tissues and/or using algorithms for the detection of potential signal peptides and, in some cases, transmembrane sequences. Notably, these approaches resulted in databases that, although largely overlapping, also had in most cases significant non-overlap in gene sets (Fig. 1A).

**Fig. 1.**
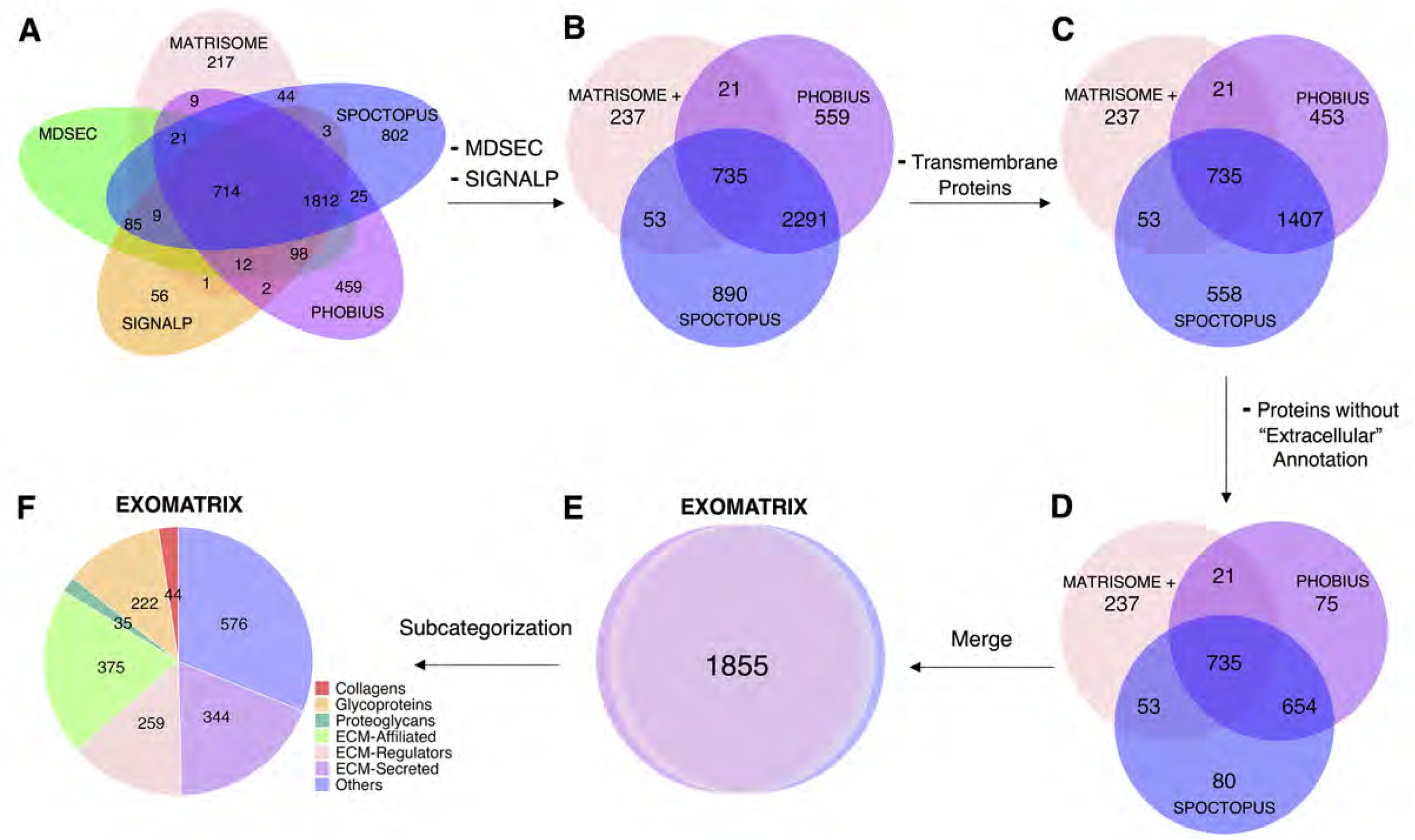
Bioinformatics pipeline used to define Exomatrix. **A)** Venn diagram showing the overlap of extracellular proteins from five databases. **B)** Venn diagram illustrating the number of proteins after the exclusion of MDSEC and manual curation of non-overlapping proteins in SIGNALP, with of 20 these proteins moved to the Matrisome dataset. **C)** Venn diagram showing the number of proteins remaining after applying a transmembrane filter. **D)** Venn diagram showing the remaining proteins after excluding those lacking the “extracellular” annotation from the Compartments resource. **E)** Diagram showing the number of proteins after merging the final three databases. **F)** Pie chart illustrating the distribution of proteins across Exomatrix subcategories (subcategories adopted and adapted from Matrisome).

To address these discrepancies, we derived from these five existing databases an integrated database of human genes that encode putative extracellular proteins. In a first step, we removed MDSEC from the Venn diagram in Fig. 1A because all its genes were also present in one or more of the other datasets. We also examined individually 56 unique genes in SignalP by performing literature searches guided by UniProt on the subcellular locations of the proteins they encode, which led to the exclusion of 36 genes that encoded solely intracellular or transmembrane proteins. The remaining 20 genes were combined with the unique Matrisome genes (Fig. 1B).

In a second step, we filtered out transmembrane proteins using the “transmembrane” versions of the databases for Spoctopus and Phobius (Fig. 1C). In a third step, using Compartments (Binder et al., 2014), a resource for subcellular localization of proteins, we filtered out proteins that did not contain the annotation term “extracellular” (Fig. 1D). Finally, we combined all the remaining genes to form a new database, Exomatrix (Fig. 1E).

Naba et al. (2012) previously provided a useful subcategorization of extracellular proteins into collagens, glycoproteins, proteoglycans, ECM-affiliated, ECM-regulators, and ECM-secreted. This previous subcategorization was adopted for the genes in Exomatrix. We manually assigned 233 new proteins to these subcategories based on UniProt and the literature. However, 576 additional genes did not clearly fit into any of Matrisome’s subcategories and were, therefore, simply included in an “others” subcategory (Fig. 1F).

### Inter-sample reproducibility of proteomic data

Given the limited availability of human and NHP fetal neocortical samples, we assessed the reproducibility of proteomic data between pairs of similarly aged samples. We used both human and NHP samples in this study under approved IACUC and IRB protocols. Given the dynamic changes in gene expression throughout cortical development, it was important to perform proteomic analysis on pairs of samples that were closest in age, namely two mid-gestation human samples at post-conceptional week (PCW) 16, and two Rhesus macaque samples at 51- and 52-days post conception (dpc), which correspond roughly to 90 dpc in humans. Samples were processed individually as described in Methods to obtain their respective proteome datasets, listed as the genes encoding the proteins detected. For each dataset, total hits were filtered to extract only the genes in Exomatrix.

For the human and NHP samples, there was a good correlation between the pairs of samples, R = 0.87 and 0.90, respectively (Fig. 2A,D) . In addition, 92% of the hits (236/277) were shared between both human samples, while 96% (220/229) were found in both NHP samples (Fig. 2B,E). A comparative list of the top 25 hits in order of abundance for each sample of a pair further illustrated these findings (Fig.2C,F), with matches present in the top 20 hits in both samples of a pair highlighted in green and matches in the top 50 in light green. Similarly, pairs of samples were also very well correlated separately for the 7 subcategories of ECM proteins described above, with few genes not appearing in both top 25 hits of paired samples (Suppl. Figs. 1 and 2). Overall, these findings support the reproducibility of the proteomic method used here.

**Fig. 2.**
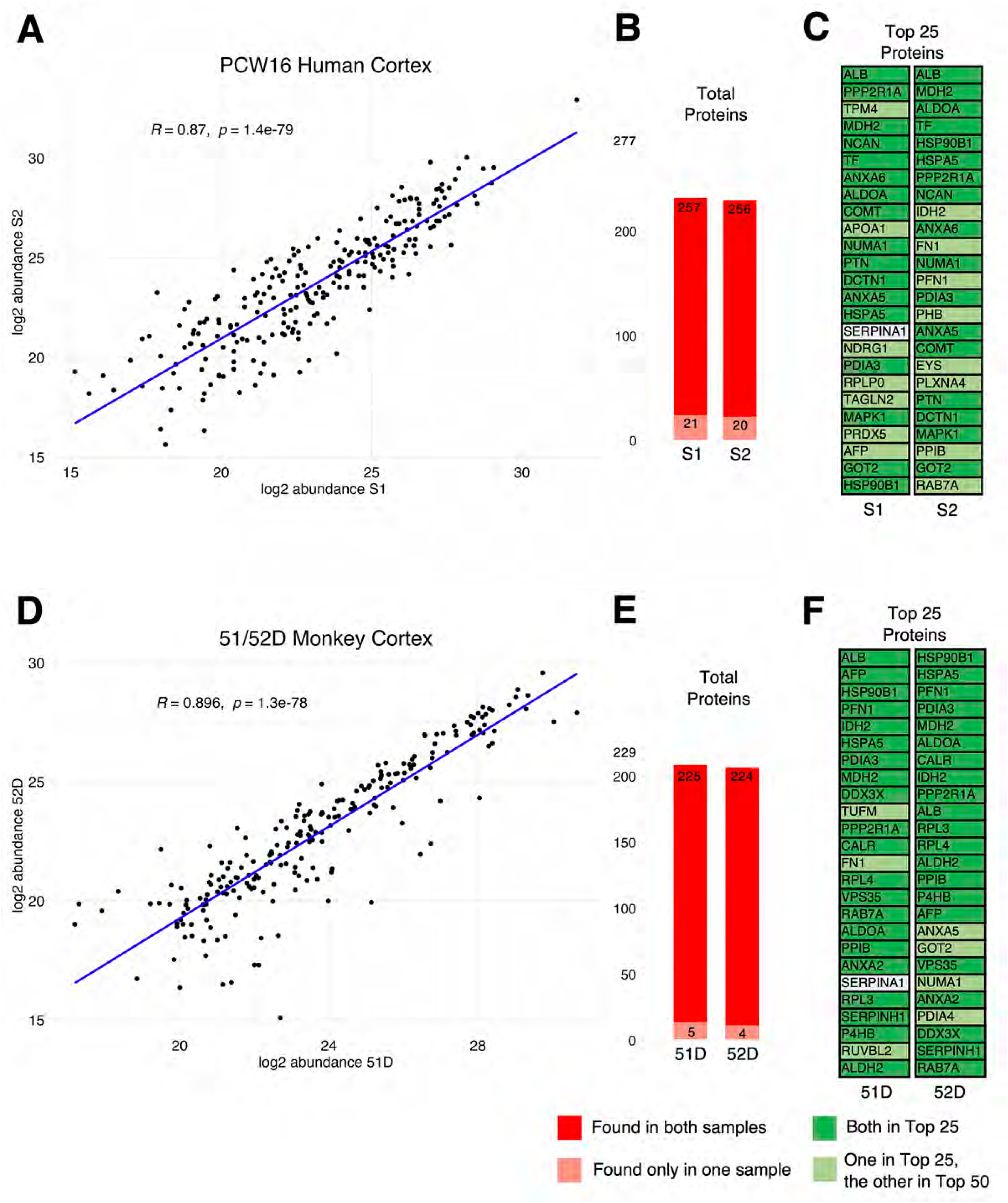
Reproducibility of proteomic data. **(A,D)** Scatter plots showing paired protein abundance between PCW16 human and 51/52D primate samples. **(B-E)** Bar plots depicting the numbers of proteins identified in each sample of a pair, with those in red in both samples and those in light red only in one. **(C-F)** Tables listing the top 25 most abundant proteins in each pair of samples.

### The extracellular milieu differs between layers in the developing cortex

To obtain more defined information about the layered distribution of ECM components in the developing neocortex, two human PCW 16 samples were microdissected into 3 layers before processing for proteomics analyses. The layers corresponded roughly to the germinal zones (a ∼650 um layer starting from the ventricular surface), the subplate and intermediate zone (the next ∼900 um thick layer), and the cortical plate and marginal zone (the outermost ∼350 um layer; n = 2; Kostovic, 2020). Surprisingly, both the proportion of the subcategories of extracellular components and the composition of each category was similar for each layer (Fig. 3A, Suppl. Fig. 3). Nevertheless, some genes for the top 25 most abundant proteins were unique to one or two layers (highlighted in orange and yellow, respectively; Fig. 3B).

**Fig. 3.**
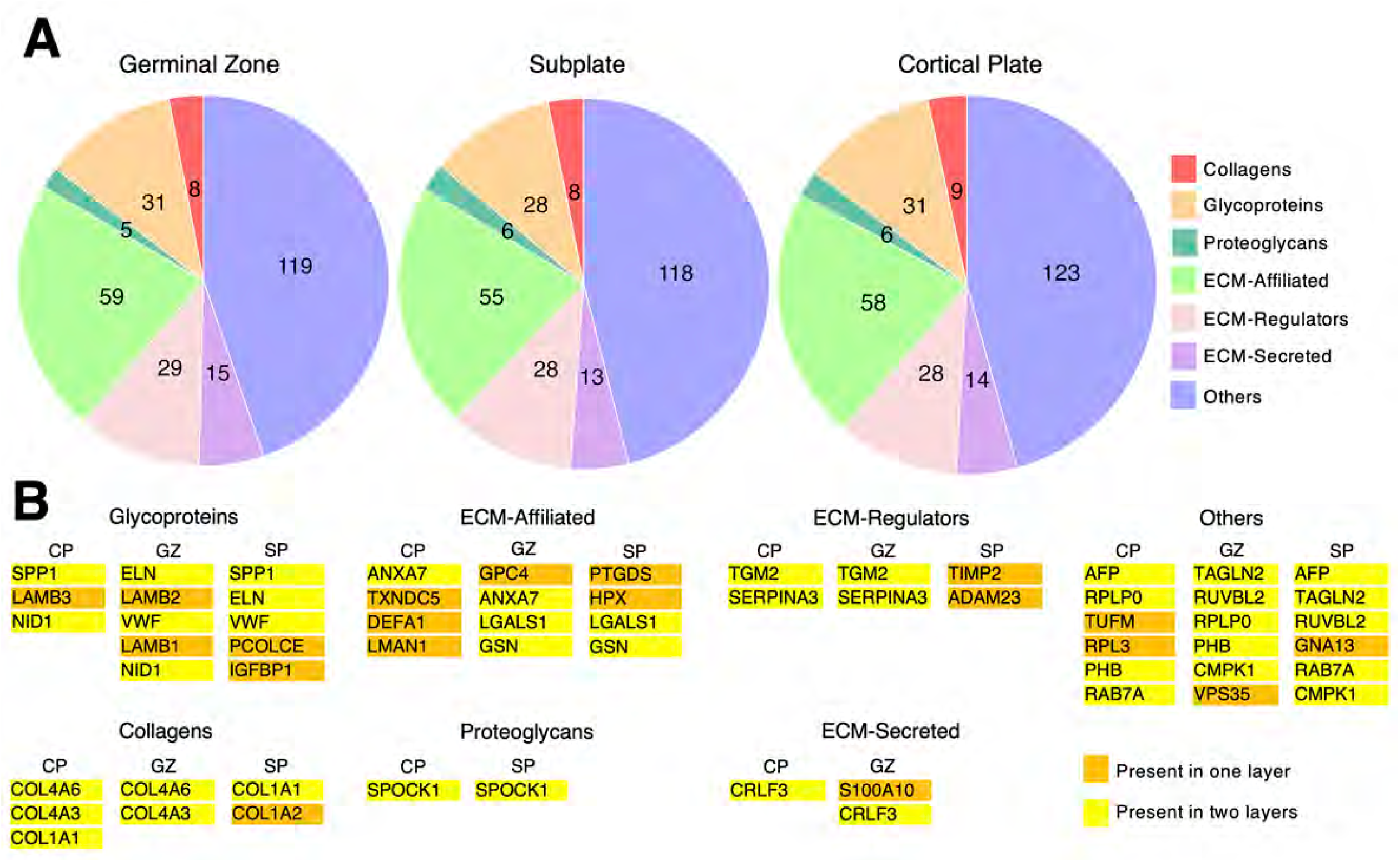
Extracellular proteins in three layers of the human developing brain. **A)** Pie charts illustrating the proportion of Exomatrix proteins by subcategory in each layer. **B)** Tables listing proteins in the top 25 that are unique to individual layers and proteins only shared by two layers.

An interesting example is osteopontin (SPP1), present in the cortical plate and subplate. This could be due to the role of SPP1 in adhesion, migration and differentiation, which are critical processes occurring in these regions as neurons establish connectivity and form mature circuits (Zhao et al. 2024). Its absence from the germinal zone could be explained by the fact that this zone is primarily focused on cell proliferation rather than migration and differentiation (Mahmud et al. 2020; Rabenstein et al. 2015). In contrast, the laminin subunits LAMB1 and LAMB2 are specific to the germinal zone, where laminins play essential roles in radial glial stem cell behavior (Radner et al. 2013).

### Comparison of the extracellular proteome and transcriptome for mid-gestational human neocortex

Proteomic and transcriptomic data from similar samples can vary in how well they correlate due to biological factors and inherent technical limitations with each approach (Haider et al., 2013). Therefore, to obtain a more reliable picture of the extracellular proteome of a tissue, it can be valuable to compare proteomic and transcriptomic datasets for similar samples. In line with this idea, we compared our Exomatrix-derived PCW 16 proteome data with published PCW 16 transcriptome data filtered for Exomatrix genes (to obtain whole tissue transcriptomes, we pooled single-cell transcriptomes from PCW 16 datasets, Polioudakis et al., 2019).

As expected, there was not a one-to-one correlation between a protein and its mRNA between samples (Fig. 4). Nevertheless, when comparing total Exomatrix proteins and mRNA based on their ranked abundance in each dataset, there was a reasonable correlation (R = 0.48, Fig. 4A). While >90% of proteins had a matching mRNA, only ∼20% of mRNAs had a matching protein (Fig. 4B), at least in part due to the larger total number of mRNA hits because of higher sensitivity for detecting mRNAs over a wider range of abundance compared with the limited range for the proteomic approach used here. The top 25 hits by abundance for the proteomic and transcriptomic datasets further reflect these differences both for the whole cortex (Fig. 4C) and by subcategory (Suppl. Fig. 4).

**Fig. 4.**
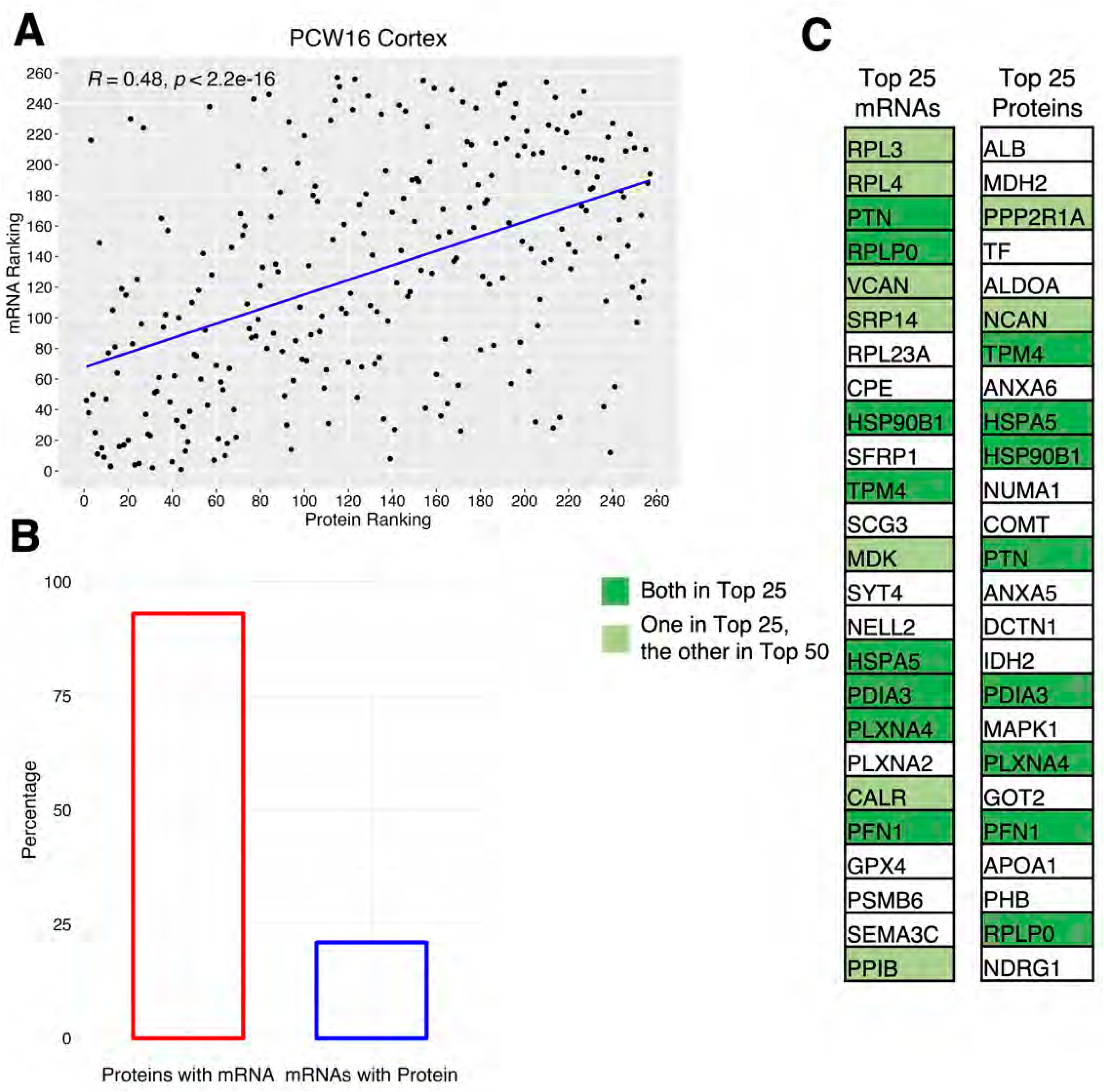
mRNA and protein correlation in PCW16 human samples. **A)** Scatter plot illustrating the ranked abundance of mRNA and protein levels for the same genes. **B)** Bar plots showing the proportion of proteins detected with their corresponding mRNA and the reverse (mRNA with corresponding proteins). **C)** Tables listing the top 25 most abundant mRNAs and proteins, highlighting the overlap.

### Comparison of the extracellular proteome and transcriptome of early human meninges

The early first trimester neocortex is a relatively simple structure comprised of a pseudostratified neuroepithelium that will generate cortical excitatory neurons and astrocytes, a marginal zone housing the earliest born Cajal-Retzius neurons, and the primitive meninges comprised of specialized fibroblast-like cells and vasculature (Meyer et al., 2000; Meyer 2007; Zecevic et al., 2010; Braun et al., 2023). While the early neuroepithelium and marginal zones have been the focus of many studies, little is known about the composition of the cell-sparse meninges.

To explore the extracellular components of the early meninges, we first performed a clustering analysis of a previously published single-cell RNA sequencing (scRNAseq) dataset for PCW 5.5 forebrain (Braun et al., 2023) (Fig. 5A; Suppl. Fig. 5). This revealed the expected cell types for this early developmental stage of brain development. Fibroblast-like cells and vascular cells, the cell types of the early meninges were identified by marker expression. All gene counts from these two cell clusters were combined (maintaining relative count abundance), filtered through Exomatrix to retain only genes encoding extracellular proteins, and grouped into the 7 extracellular subcategories.

**Fig. 5.**
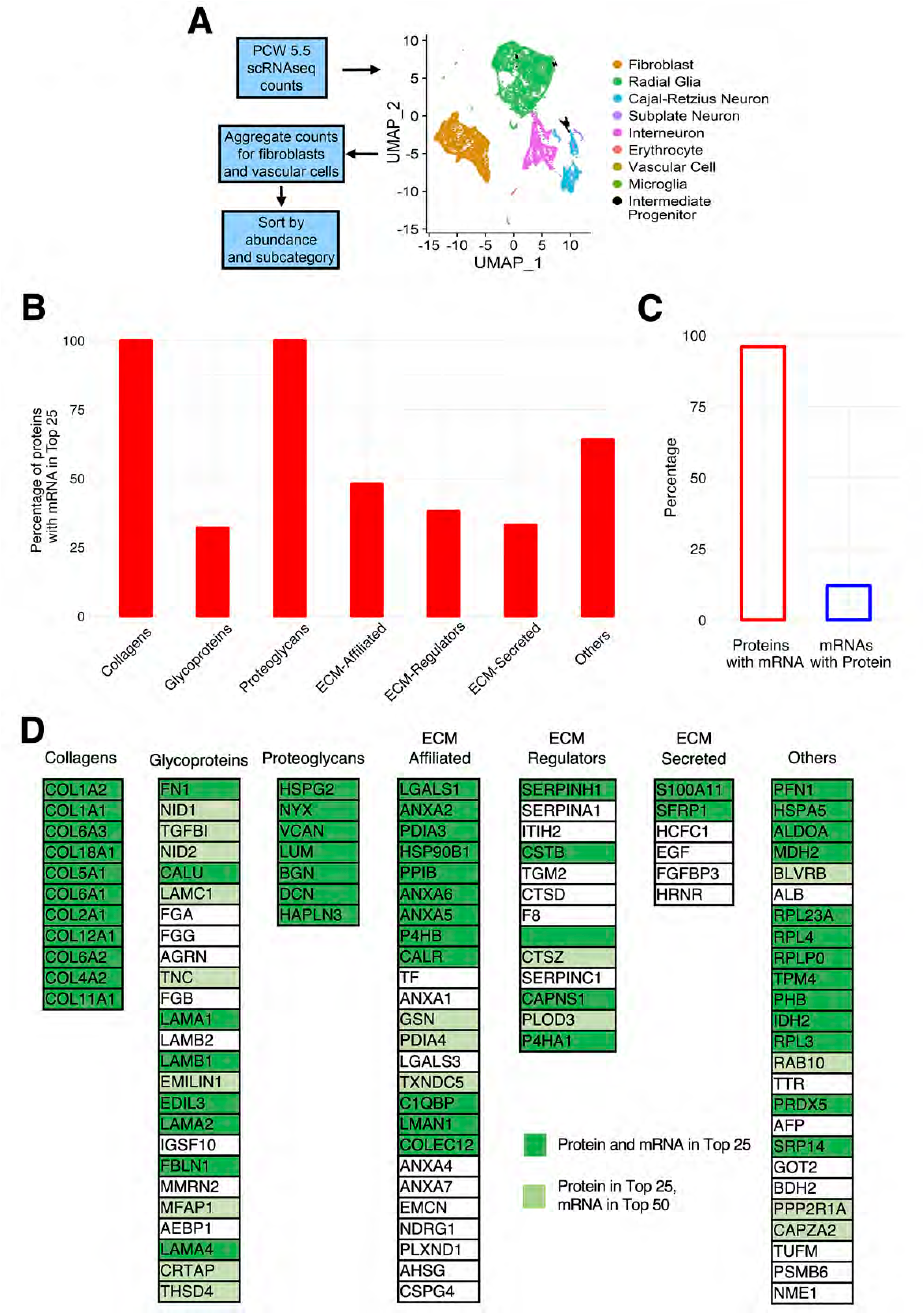
Comparison between PCW6.8 proteomic data and PCW5.5 scRNAseq data from meninges. **A)** Clustering and cell type selection for pseudo-bulk analysis. **B)** Bar plots illustrating the proportion of proteins with corresponding mRNA within each Exomatrix subcategory. **C)** Bar plots displaying the proportion of proteins with corresponding mRNAs and vice versa. **D)** Tables listing all or the top 25 most abundant proteins in each subcategory, highlighted for their corresponding mRNA abundance.

To substantiate the gene hits obtained from scRNAseq analysis, we performed proteomic analyses on a pial sample from PCW 6.8 (the best age-matched samples we were able to obtain). As with the scRNAseq data, proteomic gene hits were filtered through Exomatrix and grouped into the 7 subcategories. The most abundant hits by subcategory were then compared between scRNAseq and proteomics. The degree to which proteomic and transcriptomic datasets matched varied depending on the subcategory, but generally showed a positive correlation (Fig. 5B-D, Suppl. Fig. 6) (see also Discussions). Here, as with PCW 16 tissue, most protein hits were present in the RNA data, while the opposite was not the case (Fig. 5C), likely due to the differing sensitivity of each omics approach.

While some structural proteins identified in the early meninges, for example genes encoding Collagen 1 (COL1), Fibronectin (FN1), Nidogen 1 (NID1), Versican (VCAN), have been commonly associated with fibroblastic tissues, others were not as commonly associated, such as TGF-beta-induced protein (TGFBI), Fibrinogen alpha (FGA), IGF-binding protein 2 (IGFBP2), Nyctalopin (NYX), Osteoglycin (OGN), Osteonectin (SPOCK3), COL6, Fibulin 1 (FBLN1), Laminin (Lam111 as a heterotrimer), Perlecan (HSPG2), Lumican (LUM), Hyaluronan and proteoglycan link protein 3 (HAPLN3), and Asporin (ASPN). Some of these proteins (COL1, FN1, NID1, VCAN, TGFBI, and FGA) were confirmed by immunohistofluorescence (IHF) analyses (Fig. 6 and not shown), while others remain to be validated.

**Figure 6.**
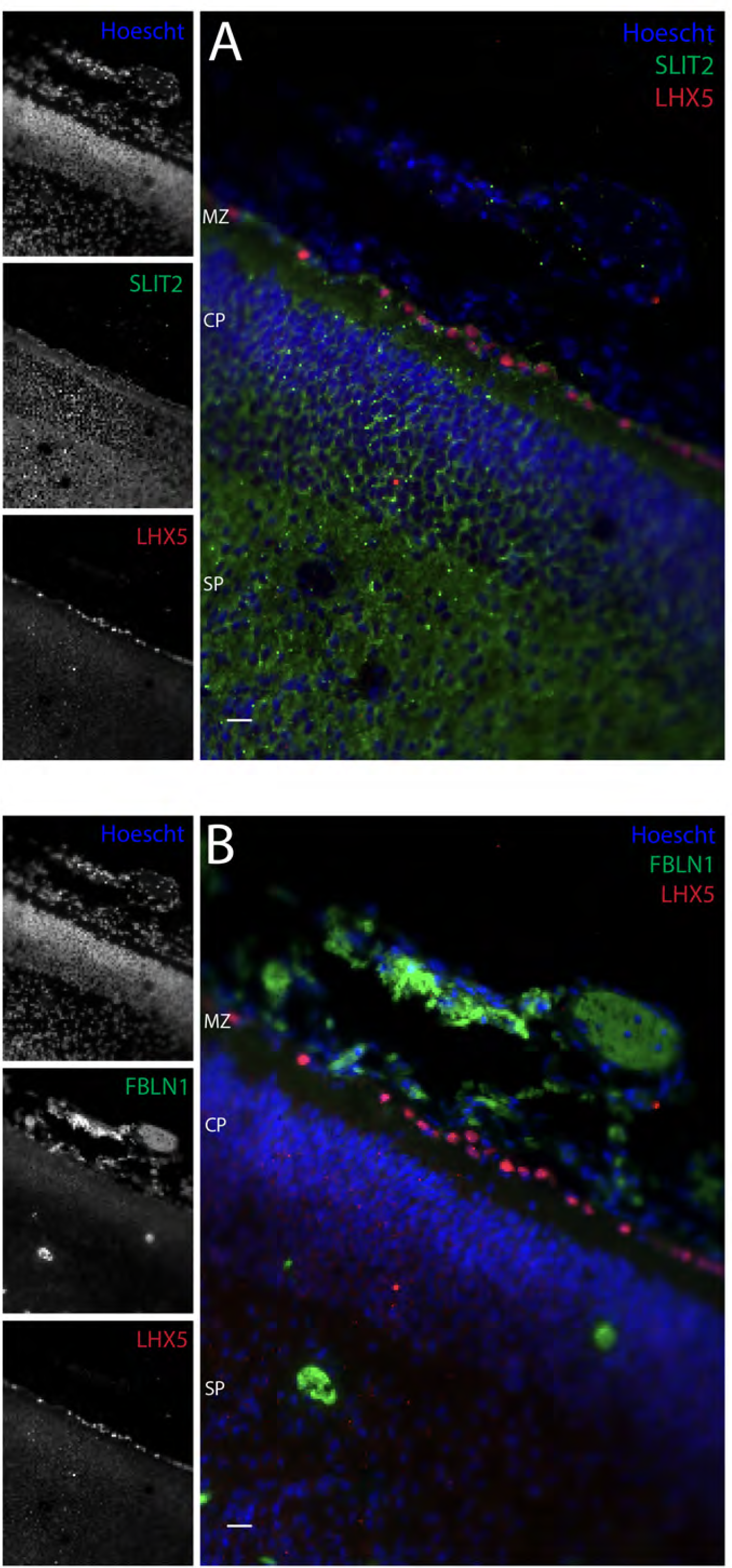
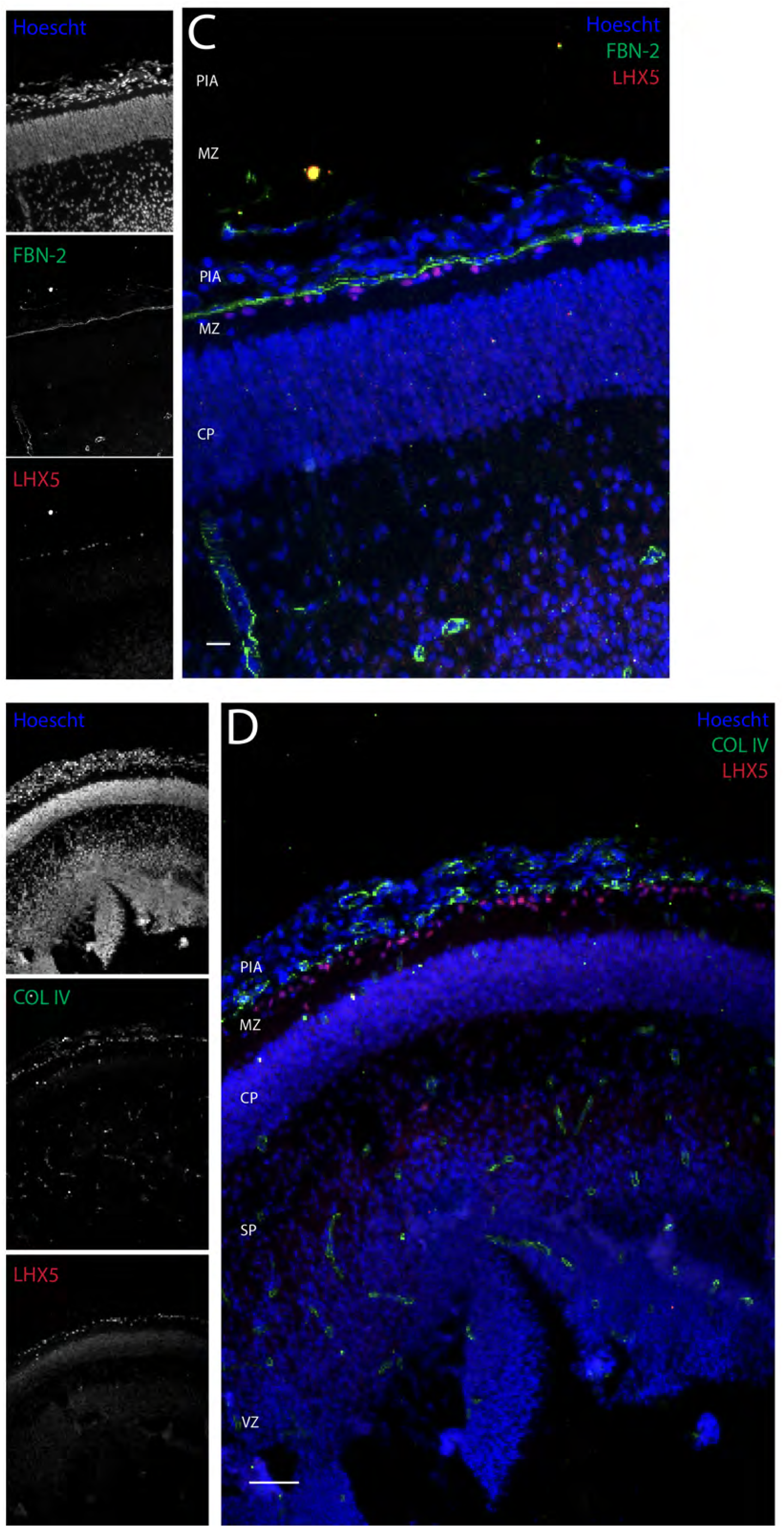
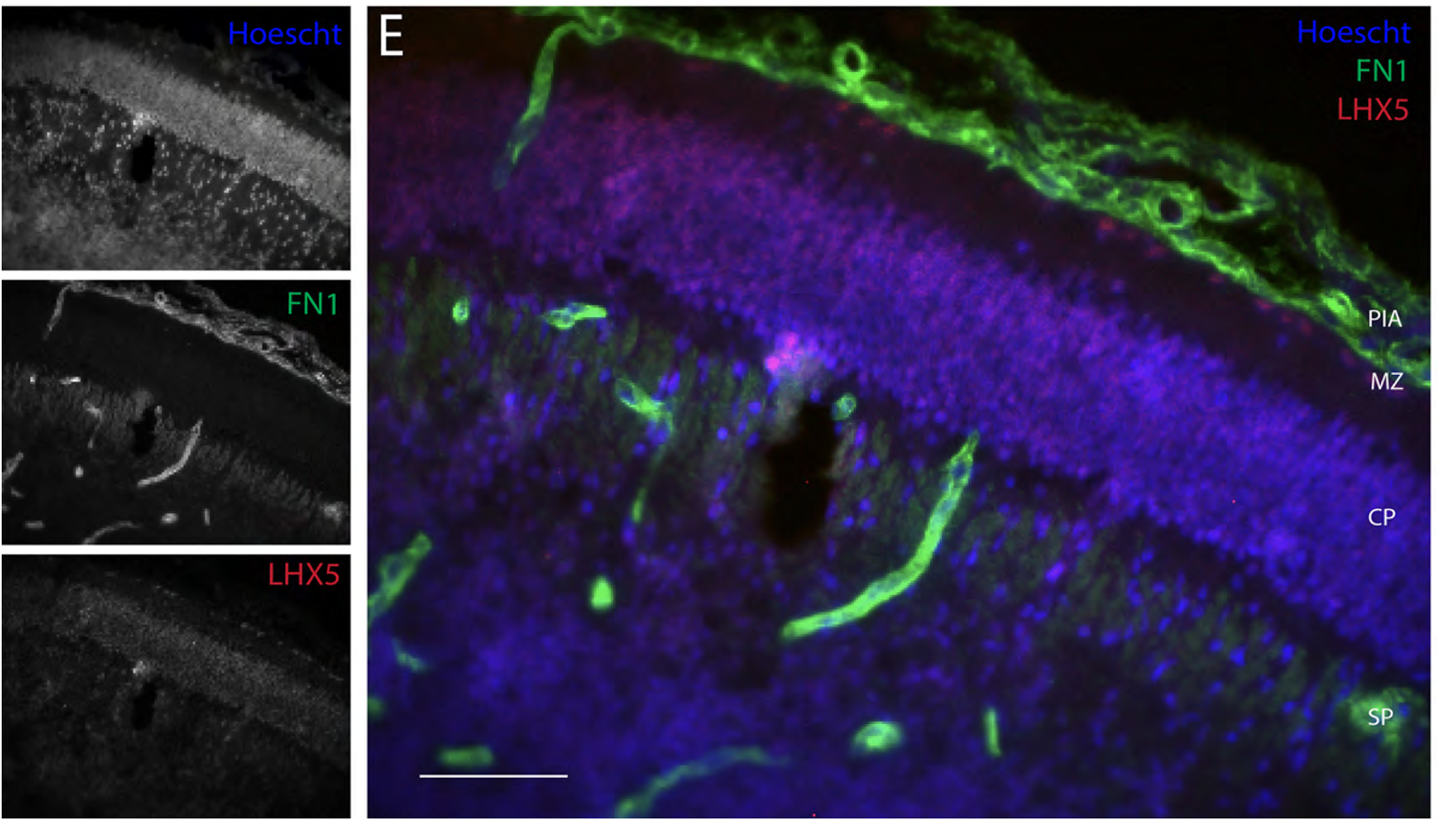
Distribution at PCW 9 of select abundant ECM proteins identified in omics analyses. LHX5-positive CRNs were used to identify the apical cortical layer and mark the separation of cortex from the developing meninges (A-E). While SLIT2 is present only within the cortical parenchyma, FBLN1 (B), COL IV (D), and FN1 (E) are observed primarily within the meninges. FBN-2 (C) exclusively marks the pial basement membrane. MZ: Marginal Zone, CP: Cortical Plate, SP: Subplate, VZ: Ventricular Zone; Scale bars: 25 μm (A, B, C), 100 μm (D, E).

Similarly, for secreted signaling and regulatory factors, some were expected such as Epidermal growth factor (EGF), CXCL12, FGF binding protein 3 (FGFBP3), Galectin 1 (LGALS1), and perhaps Semaphorin 3A (SEMA3A), while we did not anticipate others such as S100 calcium binding protein A11 (S100A11), Secreted frizzled-related protein 1 (SFRP1), Midkine (MDK), and annexins, for example Annexin A2 (ANXA2).

### Immunohistofluorescence analyses on select proteins

To learn more about the tissue distribution of certain proteins identified in the analyses above, we selected some of the most abundant hits (from either the proteomic or scRNAseq data) for IHF staining on PCW 9 sections (as we did not have sufficient PCW 6-8 tissue for IHF). In these experiments, to demarcate the developing cortical neural tissue from its overlying non-neuronal meninges, we used LHX5, which is specifically expressed in the most superficial neural cells of the developing cortex, the Cajal-Retzius neurons (CRNs) (Fig. 6A-E). Here, we present examples of IHF stains at PCW 9 that show protein distributions that are specific for different layers of the developing human neocortex.

SLIT proteins are secreted ligands known to promote axon guidance and cell migration during neural development (Gonda et al., 2020). SLIT2 was identified as a top hit, underscoring a putative role in early cortical development. SLIT2 was detected throughout the developing cortical parenchyma (including the germinal zones, not shown) and excluded from the overlying meninges (Fig. 6A).

Fibulin 1 (FBLN1) is a secreted glycoprotein that integrates into fibrillar extracellular matrix, and can interact with proteins such as laminin, nidogen, and fibrinogen (deVega et al., 2009). FBLN1 was detected throughout the developing meninges and excluded from the entire neocortical tissue proper (Fig.6B).

Decorin (DCN), FN1, and COL4 were associated with both vascular and meningeal basement membranes. And unexpectedly, Fibrilin 2 (FBN2) was present exclusively in the meningeal basement membrane separating the LHX5-positive CRNs in the marginal zone from the overlying meninges (Fig. 6C-E). Fibrilins are an understudied family of ECM proteins associated with non-neural syndromes in humans and suspected of playing essential roles in creating elasticity within tissues (Peeters et al., 2022).

## DISCUSSION

In this study, we aimed to explore the extracellular composition of the developing human and NHP neocortices using a combination of scRNAseq, proteomics, and IHF analyses. Our findings contribute to the understanding of the ECM and signaling components that guide neocortical development, offering new insights into brain formation. While we observed reasonable correlations between the proteomic and transcriptomic datasets, there were notable differences between the two. Below, we discuss this and other points, as well as their broader implications for neocortical development.

### Correlation between Proteome and Transcriptome Datasets

The correlations between proteome and transcriptome data for both the PCW 16 neocortex and meninges samples were positive but not robust. A key observation was that although most (>90%) of the detected proteins had corresponding mRNA transcripts, only 10 and 20% of the mRNA transcripts were reflected in the proteomic data. This discrepancy is likely due to several technical and biological factors, which highlight the inherent limitations of these approaches.

Proteomic methods, including those used here, tend to have lower sensitivity compared to transcriptomics. Mass spectrometry, while powerful for identifying abundant proteins, may fail to detect low-abundance proteins due to limited dynamic range. In contrast, scRNAseq can capture a broader range of mRNA transcripts, including those expressed at low levels. This likely explains why we observed many more transcripts than proteins in our datasets. In addition, not all mRNA transcripts are translated into proteins, and even when translated, protein stability and degradation rates may vary significantly, contributing to differences in detectable protein levels. Proteins with shorter half-lives may be present only transiently, making them more difficult to capture in a proteomic analysis. The slight difference in age and small sample sizes available for the proteomic and transcriptomic analyses may also contribute to the discrepancies in the presence and abundance of protein and mRNA species.

Certain proteins detected in the ECM proteome may come from external sources, especially relevant for analysis of the meninges, which are highly vascularized and in which the CSF is beginning to form. For example, circulating or diffusible factors may be detected that are not synthesized by the local cells. This would lead to the detection of these proteins in the proteome but not in the transcriptome of the source tissue, as its cells may not be actively transcribing the corresponding genes.

Lastly, technical limitations of both methods could contribute to the discrepancies. For example, scRNAseq, while highly sensitive, has its own biases, particularly in capturing transcripts from certain cell types or cellular compartments. Variability in sample processing, such as the dissection and preservation of tissues, could also influence the composition of detected mRNAs and proteins. Together, these factors likely contribute to the moderate correlation between the proteomic and transcriptomic datasets, especially for complex tissues like the developing neocortex and meninges.

### The similarity between ECM Gene Sets across Cortical Layers at PCW 16

Another notable finding was the similarity in ECM gene sets across the cortical plate, subplate, and germinal zones at PCW 16, despite the expected differences in the cellular composition of these layers. This similarity may be partly explained by the structural organization of the developing cortex and the imprecision of the dissections.

The cortical plate and germinal zones are cell-dense regions with relatively little ECM. These zones primarily comprise precursor cells, proliferating neuroblasts, and differentiating neurons. In contrast, the subplate is a large, cell-sparse region rich in ECM. This region serves as a critical scaffold during development, guiding the migration of neurons and the establishment of cortical circuits.

It is also possible that the process of microdissection, imprecise at best, that was used for proteomic analysis introduced some contamination between layers. Namely, the subplate’s extensive ECM could have been inadvertently sampled during the dissection of the cortical plate and germinal zone tissues, leading to an overlap in detected ECM components. Even with careful microdissection, the fine structural boundaries between these layers are difficult to preserve, particularly in the very soft developing brain, where layers are not yet fully differentiated.

In addition, or alternatively, the overlap in ECM composition might reflect shared developmental cues that influence multiple cortical layers. Structural and signaling components, such as glycoproteins and growth factors, may diffuse across cortical layers, playing roles in cell differentiation and tissue organization. Thus, the similarity in ECM composition could indicate common extracellular signals regulating neocortical development at this stage despite the cell differences between layers. Further studies, such as IHF and RNA in situ hybridization analyses, will be required to shed more light on these possibilities.

### Broader Implications

The findings presented in this study offer several important implications for the understanding of neocortical development. First, the moderate correlation between proteome and transcriptome datasets highlights the importance of employing complementary omics approaches to fully capture the molecular landscape of developing tissues. While transcriptomics provides a detailed map of gene expression, proteomics offers a more functional readout of the proteins actively shaping tissue development. Together, these methods can provide a more comprehensive view of the dynamic interactions between cells and their extracellular environment.

Second, the observed similarities in ECM composition across cortical layers underscore the need for careful consideration of both tissue architecture and experimental methodology in developmental studies. The ECM plays a critical role in guiding neural precursor cells, and even subtle variations in ECM composition between layers may have significant effects on cortical organization and function. Future studies aimed at dissecting the specific roles of some of the ECM components identified here will be crucial for understanding how these extracellular elements contribute to normal neocortex development and the etiology of neurodevelopmental disorders.

In conclusion, this study provides valuable insights into the extracellular composition of the developing neocortex and meninges. The integration of transcriptomic and proteomic data has illuminated new avenues for hypothesis-driven research, which could lead to a deeper understanding of the molecular mechanisms that shape cortical development and underlie neurodevelopmental diseases.

## METHODS

### Sample sources

Specimens from PCW 5.5-16 human cortices were obtained from the Human Development Biology Resource at the University College London and Newcastle University, the Birth Defects Research Laboratory at the University of Washington, and the Jack D. Weiler Hospital, Montefiore, with ethics board approval and maternal written consent. This study was performed in accordance with ethical and legal guidelines of the Albert Einstein College of Medicine institutional review board. Time-mated breeding of rhesus macaque males and females was performed to obtain fetal samples and relatively specific days after conception, which was accomplished by measuring estradiol daily in the female starting from D5 to D8 after menses began (Sosa et al., 2018). Pregnancy was confirmed by measuring progesterone as well as by ultrasound. Fetuses at 51 and 52 dpc were collected by C-section. All rhesus macaque time-mated breeding experiments were conducted following the approval of the Oregon National Primate Research Center (ONPRC) Institutional Animal Care and Use Committee (IACUC).

### In Silico definition of Exomatrix

The Matrisome database (Naba et al., 2012) was obtained through the GSEA (Gene Set Enrichment Analysis) and MSigDB (Molecular Signatures Database) online resources, accessible at: https://www.gsea-msigdb.org/gsea/msigdb/index.jsp. Data from SIGNALP, MDSEC, SPOCTOPUS, and PHOBIUS extracellular and membrane databases were retrieved through the Human Protein Atlas repository (Uhlén et al., 2015), available at https://www.proteinatlas.org. The COMPARTMENTS database (Binder et al., 2014) was utilized to gather and visualize protein subcellular localization evidence, accessible at https://compartments.jensenlab.org. Manual curation of the protein list was guided by the UniProt database (UniProt Consortium, 2023), available at https://www.uniprot.org. All data filtering steps were conducted in R using the VennDiagram, dplyr, and writexl packages. The code for this analysis is available at: https://github.com/fvilicich/omicspaper/blob/main/OmicsPaper.

### mRNA and protein comparisons

Protein-protein and protein-mRNA correlations in human and primate samples were analyzed using the built-in stats package in R. When comparing mRNA vs. protein data, both datasets were ranked by abundance and then compared using Spearman’s coefficient. Statistical significance was set at p < 0.05. Scatter and bar plots were illustrated with the ggplot2 package in R.

### Sample preparation for proteomics

Samples were flash-frozen in liquid nitrogen and stored at -80°C until the extraction day. Upon extraction, 2% SDS extraction buffer was added (2% SDS, 20 mM TEAB, 2 mM DTT), and the sample was sonicated in 3-second bursts until homogenization was achieved. The homogenate was centrifuged at 12,000 RPM for 30 seconds. The supernatant was collected as Fraction A, while the pellet was resuspended in 5% SDS extraction buffer (5% SDS, 50 mM TEAB, 5 mM DTT). The sample was centrifuged again at 12,000 RPM for 30 seconds, and the supernatant was collected as Fraction B. Both fractions (A and B) were combined and submitted for mass spectrometry analysis.

### Pseudobulk of Pial Fibroblasts and Vascular Cells

Using the count tables from Braun et al. (2023) for PCW 5.5 forebrain scRNAseq, we identified cell clusters using Seurat v4 with the default clustering parameters in the PCA space. Using Feature plots and differentially expressed gene (DEG) lists identified by the Wilcox Rank Sum Test of each cluster to all other clusters (see Supplementary Table 1), we annotated each cluster by cell type based on the expression of canonical forebrain cell type markers. After identifying cell types, we subset the pial fibroblast-like and vascular cells and aggregated the counts of the cells belonging to this subset for each gene. Code is available at https://github.com/Sesukai87/Omics-Paper/blob/main/Omics_Paper_Code.R. Next, we selected only those genes that fell into the 7 ECM subcategory groups from Exomatrix. Finally, We identified the most abundant ECM genes in each pseudo-layer based on the aggregate counts in each subcategory).

### Immunohistofluorescence

Formalin-fixed paraffin-embedded (FFPE) blocks of cortical tissue, aged PCW 9, were provided by the Human Development Biology Resource center (London, United Kingdom). Blocks were processed at the Histology & Comparative Pathology Core at Albert Einstein College of Medicine (Bronx, NY, USA) and sectioned at 25 μm. The sections were deparaffinized and rehydrated by undergoing the following washes: xylene (3 min) twice, xylene:ethanol (3 min) twice, 100% ethanol (5 min), 95% ethanol (3 min), 75% ethanol (3 min), 50% ethanol (3 min), water (3 min) twice. Antigen retrieval was performed on all sections by submerging slides in sodium citrate buffer (10mM sodium citrate, 0.05% Tween-20, pH 6.0) and heating in a microwave at 98°C for 20 minutes. Sections were incubated in 50 mM glycine at RT on a shaker for 5 minutes and washed with 1X PBS for 5 minutes. Primary antibodies (see Table immediately below) were prepared in blocking buffer (5% donkey serum and 0.025% sodium azide in 0.3% Triton X-100 in DPBS) and added to each section for incubation at 4°C overnight. The sections were washed twice with PBS for 5 minutes and incubated with secondary antibody (1:500) combinations for 2 hours on a shaker at RT and Hoescht for 5 minutes at RT. Each slide was washed with PBS twice for 5 minutes, mounted with Flouromount-G (ThermoFisher, New Jersey, United States, cat. 00-4958-02) and covered with a glass coverslip. Sections were imaged with an epifluorescence microscope and analyzed using Adobe Photoshop.

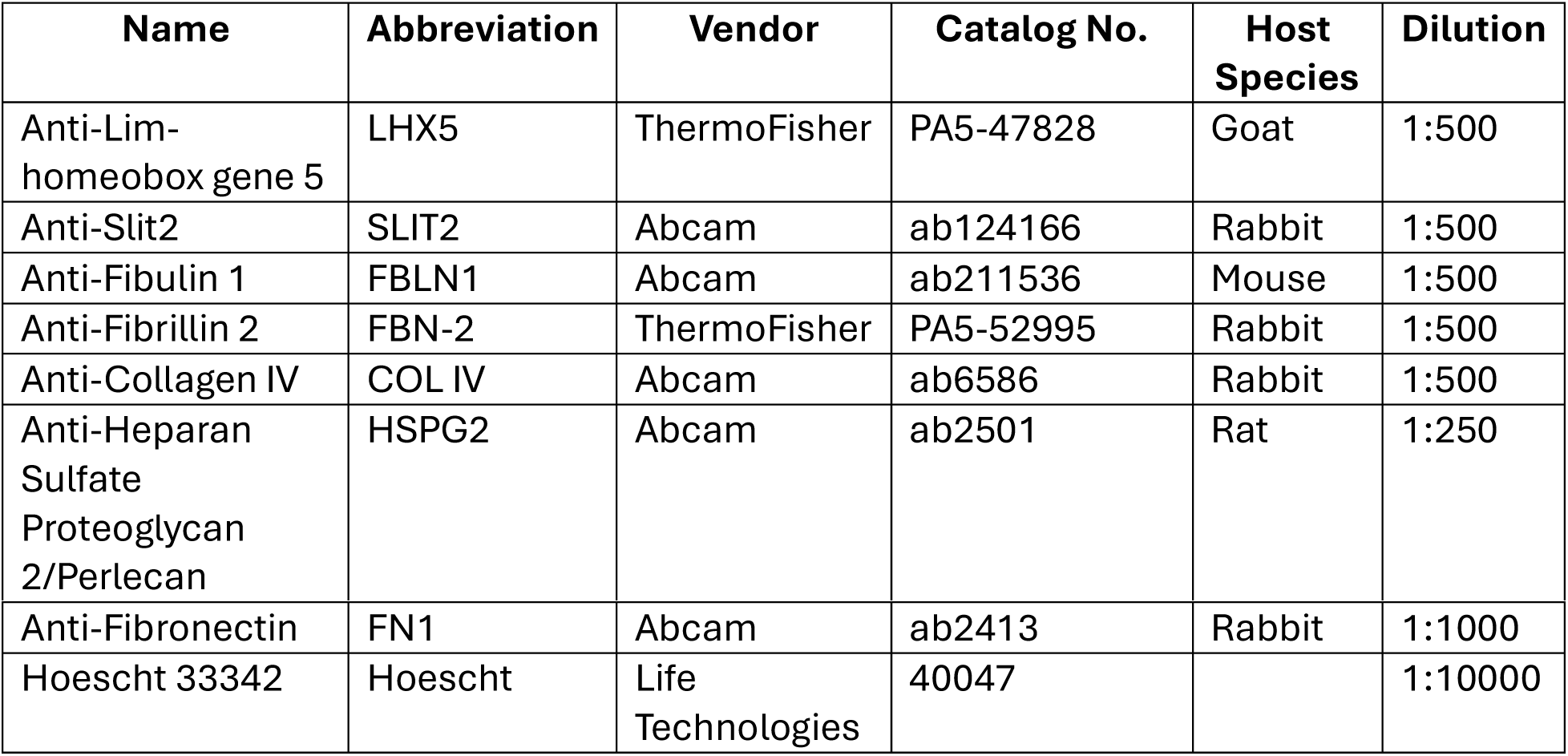

| **Name** | **Abbreviation** | **Vendor** | **Catalog No.** | **Host Species** | **Dilution** |
| --- | --- | --- | --- | --- | --- |
| Anti-Lim-homeobox gene 5 | LHX5 | ThermoFisher | PA5-47828 | Goat | 1:500 |
| Anti-Slit2 | SLIT2 | Abcam | ab124166 | Rabbit | 1:500 |
| Anti-Fibulin 1 | FBLN1 | Abcam | ab211536 | Mouse | 1:500 |
| Anti-Fibrillin 2 | FBN-2 | ThermoFisher | PA5-52995 | Rabbit | 1:500 |
| Anti-Collagen IV | COL IV | Abcam | ab6586 | Rabbit | 1:500 |
| Anti-Heparan Sulfate<br>Proteoglycan 2/Perlecan | HSPG2 | Abcam | ab2501 | Rat | 1:250 |
| Anti-Fibronectin | FN1 | Abcam | ab2413 | Rabbit | 1:1000 |
| Hoescht 33342 | Hoescht | Life Technologies | 40047 |  | 1:10000 |

## AUTHOR CONTRIBUTIONS

FV lead the development of Exomatrix and the subcategorization of its genes, prepared samples for proteomics, performed proteome-transcriptome comparisons, and contributed to manuscript preparation. DV performed hand curation of genes for exclusion from the database and prepared samples for proteomic analyses. SKG converted raw data to usable formats and performed pseudo-layering from published datasets. NN performed hand curation for Exomatrix and performed IHF experiments. NP performed IHF experiments. AQ performed hand curation of Exomatrix dataset. NB, EG, ES, HN, SP, JA, NM, NS, AJ, SL, and IG provided samples. MM and JH provided NHP samples. SS performed proteomics analyses.

## ACKNOWLEDGEMENTS

The human embryonic and fetal material was provided in part by the Joint MRC / Wellcome (MR/R006237/1, MR/X008304/1, and 226202/Z/22/Z) Human Developmental Biology Resource (www.hdbr.org). Funding was provided by NIH under NICHD Grant # R24HD000836, to I.A.G. and to J.H under OD Grant # P51 OD011092. We acknowledge Kimberly A. Aldinger, Dan Doherty, Ian G. Phelps, Jennifer C. Dempsey, Mei Deng, Eric Y. So, Yasmeen Otaibi, Lucinda A. Cort for their support in sample processing. We also acknowledge the P51 OD011092 supported ONPRC Pathology Services Unit, Surgical Services Unit, Endocrine Technology Core, and the expert animal care staff in the Division of Comparative Medicine for their help and support of the NHP studies.

## Notes

### Competing Interest Statement

The authors have declared no competing interest.

## References

1. Binder JX, Pletscher-Frankild S, Tsafou K, Stolte C, O’Donoghue SI, Schneider R, Jensen LJ. COMPARTMENTS: unification and visualization of protein subcellular localization evidence. Database (Oxford). 2014 Feb 25;2014:bau012. doi: 10.1093/database/bau012. PMID: 24573882; PMCID: PMC3935310.

2. Braun E, Danan-Gotthold M, Borm LE, Lee KW, Vinsland E, Lönnerberg P, Hu L, Li X, He X, Andrusivová Ž, Lundeberg J, Barker RA, Arenas E, Sundström E, Linnarsson S. Comprehensive cell atlas of the first-trimester developing human brain. Science. 2023 Oct 13;382(6667):eadf1226. doi: 10.1126/science.adf1226. Epub 2023 Oct 13. PMID: 37824650.

3. DeFelipe J, López-Cruz PL, Benavides-Piccione R, Bielza C, Larrañaga P, Anderson S, Burkhalter A, Cauli B, Fairén A, Feldmeyer D, Fishell G, Fitzpatrick D, Freund TF, González-Burgos G, Hestrin S, Hill S, Hof PR, Huang J, Jones EG, Kawaguchi Y, Kisvárday Z, Kubota Y, Lewis DA, Marín O, Markram H, McBain CJ, Meyer HS, Monyer H, Nelson SB, Rockland K, Rossier J, Rubenstein JL, Rudy B, Scanziani M, Shepherd GM, Sherwood CC, Staiger JF, Tamás G, Thomson A, Wang Y, Yuste R, Ascoli GA. New insights into the classification and nomenclature of cortical GABAergic interneurons. Nat Rev Neurosci. 2013 Mar;14(3):202–16. doi: 10.1038/nrn3444. Epub 2013 Feb 6. PMID: 23385869; PMCID: PMC3619199.

4. de Vega S, Iwamoto T, Yamada Y. Fibulins: multiple roles in matrix structures and tissue functions. Cell Mol Life Sci. 2009 Jun;66(11-12):1890–902. doi: 10.1007/s00018-009-8632-6. PMID: 19189051; PMCID: PMC11115505.

5. Eze UC, Bhaduri A, Haeussler M, Nowakowski TJ, Kriegstein AR. Single-cell atlas of early human brain development highlights heterogeneity of human neuroepithelial cells and early radial glia. Nat Neurosci. 2021 Apr;24(4):584–594. doi: 10.1038/s41593-020-00794-1. Epub 2021 Mar 15. PMID: 33723434; PMCID: PMC8012207.

6. Gonda Y, Namba T, Hanashima C. Beyond Axon Guidance: Roles of Slit-Robo Signaling in Neocortical Formation. Front Cell Dev Biol. 2020 Dec 23;8:607415. doi: 10.3389/fcell.2020.607415. PMID: 33425915; PMCID: PMC7785817.

7. Greig LC, Woodworth MB, Galazo MJ, Padmanabhan H, Macklis JD. Molecular logic of neocortical projection neuron specification, development and diversity. Nat Rev Neurosci. 2013 Nov;14(11):755–69. doi: 10.1038/nrn3586. Epub 2013 Oct 9. PMID: 24105342; PMCID: PMC3876965.

8. Grove EA, Fukuchi-Shimogori T. Generating the cerebral cortical area map. Annu Rev Neurosci. 2003;26:355–80. doi: 10.1146/annurev.neuro.26.041002.131137. PMID: 14527269.

9. Haider S, Pal R. Integrated analysis of transcriptomic and proteomic data. Curr Genomics. 2013 Apr;14(2):91–110. doi: 10.2174/1389202911314020003. PMID: 24082820; PMCID: PMC3637682.

10. Hao Y, Hao S, Andersen-Nissen E, Mauck WM 3rd, Zheng S, Butler A, Lee MJ, Wilk AJ, Darby C, Zager M, Hoffman P, Stoeckius M, Papalexi E, Mimitou EP, Jain J, Srivastava A, Stuart T, Fleming LM, Yeung B, Rogers AJ, McElrath JM, Blish CA, Gottardo R, Smibert P, Satija R. Integrated analysis of multimodal single-cell data. Cell. 2021 Jun 24;184(13):3573–3587.e29. doi: 10.1016/j.cell.2021.04.048. Epub 2021 May 31. PMID: 34062119; PMCID: PMC8238499.

11. Käll L, Krogh A, Sonnhammer EL. A combined transmembrane topology and signal peptide prediction method. J Mol Biol. 2004 May 14;338(5):1027–36. doi: 10.1016/j.jmb.2004.03.016. PMID: 15111065.

12. Kostović I. The enigmatic fetal subplate compartment forms an early tangential cortical nexus and provides the framework for construction of cortical connectivity. Prog Neurobiol. 2020 Nov;194:101883. doi: 10.1016/j.pneurobio.2020.101883. Epub 2020 Jul 11. PMID: 32659318.

13. Leone DP, Srinivasan K, Chen B, Alcamo E, McConnell SK. The determination of projection neuron identity in the developing cerebral cortex. Curr Opin Neurobiol. 2008 Feb;18(1):28–35. doi: 10.1016/j.conb.2008.05.006. Epub 2008 May 26. PMID: 18508260; PMCID: PMC2483251.

14. Mahmud FJ, Du Y, Greif E, Boucher T, Dannals RF, Mathews WB, Pomper MG, Sysa-Shah P, Metcalf Pate KA, Lyons C, Carlson B, Chacona M, Brown AM. Osteopontin/secreted phosphoprotein-1 behaves as a molecular brake regulating the neuroinflammatory response to chronic viral infection. J Neuroinflammation. 2020 Sep 17;17(1):273. doi: 10.1186/s12974-020-01949-4.

15. McConnell SK. Constructing the cerebral cortex: neurogenesis and fate determination. Neuron. 1995 Oct;15(4):761–8. doi: 10.1016/0896-6273(95)90168-x. PMID: 7576626.

16. Meyer G, Schaaps JP, Moreau L, Goffinet AM. Embryonic and early fetal development of the human neocortex. J Neurosci. 2000 Mar 1;20(5):1858–68. doi: 10.1523/JNEUROSCI.20-05-01858.2000. PMID: 10684887; PMCID: PMC6772901.

17. Meyer G. Genetic control of neuronal migrations in human cortical development. Adv Anat Embryol Cell Biol. 2007;189:1 p preceding 1, 1-111. PMID: 17212070.

18. Naba A, Clauser KR, Hoersch S, Liu H, Carr SA, Hynes RO. The matrisome: in silico definition and in vivo characterization by proteomics of normal and tumor extracellular matrices. Mol Cell Proteomics. 2012 Apr;11(4):M111.014647. doi: 10.1074/mcp.M111.014647. Epub 2011 Dec 9. PMID: 22159717; PMCID: PMC3322572.

19. O’Leary DD, Chou SJ, Sahara S. Area patterning of the mammalian cortex. Neuron. 2007 Oct 25;56(2):252–69. doi: 10.1016/j.neuron.2007.10.010. PMID: 17964244.

20. Peeters S, De Kinderen P, Meester JAN, Verstraeten A, Loeys BL. The fibrillinopathies: New insights with focus on the paradigm of opposing phenotypes for both FBN1 and FBN2. Hum Mutat. 2022 Jul;43(7):815–831. doi: 10.1002/humu.24383. Epub 2022 Apr 28. PMID: 35419902; PMCID: PMC9322447.

21. Polioudakis D, de la Torre-Ubieta L, Langerman J, Elkins AG, Shi X, Stein JL, Vuong CK, Nichterwitz S, Gevorgian M, Opland CK, Lu D, Connell W, Ruzzo EK, Lowe JK, Hadzic T, Hinz FI, Sabri S, Lowry WE, Gerstein MB, Plath K, Geschwind DH. A Single-Cell Transcriptomic Atlas of Human Neocortical Development during Mid-gestation. Neuron. 2019 Sep 4;103(5):785–801.e8. doi: 10.1016/j.neuron.2019.06.011. Epub 2019 Jul 11. PMID: 31303374; PMCID: PMC6831089.

22. Rabenstein M, Hucklenbroich J, Willuweit A, Ladwig A, Fink GR, Schroeter M, Langen KJ, Rueger MA. Osteopontin mediates survival, proliferation and migration of neural stem cells through the chemokine receptor CXCR4. Stem Cell Res Ther. 2015 May 22;6(1):99. doi: 10.1186/s13287-015-0098-x. PMID: 25998490; PMCID: PMC4464234.

23. Radner S, Banos C, Bachay G, Li YN, Hunter DD, Brunken WJ, Yee KT. β2 and γ3 laminins are critical cortical basement membrane components: ablation of Lamb2 and Lamc3 genes disrupts cortical lamination and produces dysplasia. Dev Neurobiol. 2013 Mar;73(3):209–29. doi: 10.1002/dneu.22057. Epub 2012 Oct 25. PMID: 22961762.

24. Sosa E, Chen D, Rojas EJ, Hennebold JD, Peters KA, Wu Z, Lam TN, Mitchell JM, Sukhwani M, Tailor RC, Meistrich ML, Orwig KE, Shetty G, Clark AT. Differentiation of primate primordial germ cell-like cells following transplantation into the adult gonadal niche. Nat Commun. 2018 Dec 17;9(1):5339. doi: 10.1038/s41467-018-07740-7. PMID: 30559363; PMCID: PMC6297357.

25. Teufel F, Almagro Armenteros JJ, Johansen AR, Gíslason MH, Pihl SI, Tsirigos KD, Winther O, Brunak S, von Heijne G, Nielsen H. SignalP 6.0 predicts all five types of signal peptides using protein language models. Nat Biotechnol. 2022 Jul;40(7):1023–1025. doi: 10.1038/s41587-021-01156-3. Epub 2022 Jan 3. PMID: 34980915; PMCID: PMC9287161.

26. Uhlén M, Fagerberg L, Hallström BM, Lindskog C, Oksvold P, Mardinoglu A, Sivertsson Å, Kampf C, Sjöstedt E, Asplund A, Olsson I, Edlund K, Lundberg E, Navani S, Szigyarto CA, Odeberg J, Djureinovic D, Takanen JO, Hober S, Alm T, Edqvist PH, Berling H, Tegel H, Mulder J, Rockberg J, Nilsson P, Schwenk JM, Hamsten M, von Feilitzen K, Forsberg M, Persson L, Johansson F, Zwahlen M, von Heijne G, Nielsen J, Pontén F. Proteomics. Tissue-based map of the human proteome. Science. 2015 Jan 23;347(6220):1260419. doi: 10.1126/science.1260419. PMID: 25613900.

27. UniProt Consortium. UniProt: the Universal Protein Knowledgebase in 2023. Nucleic Acids Res. 2023 Jan 6;51(D1):D523–D531. doi: 10.1093/nar/gkac1052. PMID: 36408920; PMCID: PMC9825514.

28. Viklund H, Bernsel A, Skwark M, Elofsson A. SPOCTOPUS: a combined predictor of signal peptides and membrane protein topology. Bioinformatics. 2008 Dec 15;24(24):2928–9. doi: 10.1093/bioinformatics/btn550. Epub 2008 Oct 22. PMID: 18945683.

29. Wang L, Wang C, Moriano JA, Chen S, Zuo G, Cebrián-Silla A, Zhang S, Mukhtar T, Wang S, Song M, de Oliveira LG, Bi Q, Augustin JJ, Ge X, Paredes MF, Huang EJ, Alvarez-Buylla A, Duan X, Li J, Kriegstein AR. Molecular and cellular dynamics of the developing human neocortex at single-cell resolution. bioRxiv [Preprint]. 2024 Aug 4:2024.01.16.575956. doi: 10.1101/2024.01.16.575956. PMID: 39131371; PMCID: PMC11312442.

30. Zecevic N, Hu F, Jakovcevski I. Interneurons in the developing human neocortex. Dev Neurobiol. 2011 Jan 1;71(1):18–33. doi: 10.1002/dneu.20812. PMID: 21154907; PMCID: PMC3117059.

31. Zeng B, Liu Z, Lu Y, Zhong S, Qin S, Huang L, Zeng Y, Li Z, Dong H, Shi Y, Yang J, Dai Y, Ma Q, Sun L, Bian L, Han D, Chen Y, Qiu X, Wang W, Marín O, Wu Q, Wang Y, Wang X. The single-cell and spatial transcriptional landscape of human gastrulation and early brain development. Cell Stem Cell. 2023 Jun 1;30(6):851–866.e7. doi: 10.1016/j.stem.2023.04.016. Epub 2023 May 15. PMID: 37192616; PMCID: PMC10241223.

32. Zhao Y, Huang Z, Gao L, Ma H, Chang R. Osteopontin/SPP1: a potential mediator between immune cells and vascular calcification. Front Immunol. 2024 Jun 11;15:1395596. doi: 10.3389/fimmu.2024.1395596. PMID: 38919629; PMCID: PMC11196619.

