## Supplemental Figure 1-6 for "Multi-omics identification of extracellular components of the fetal monkey and human neocortex"

Suppl. Fig. 1

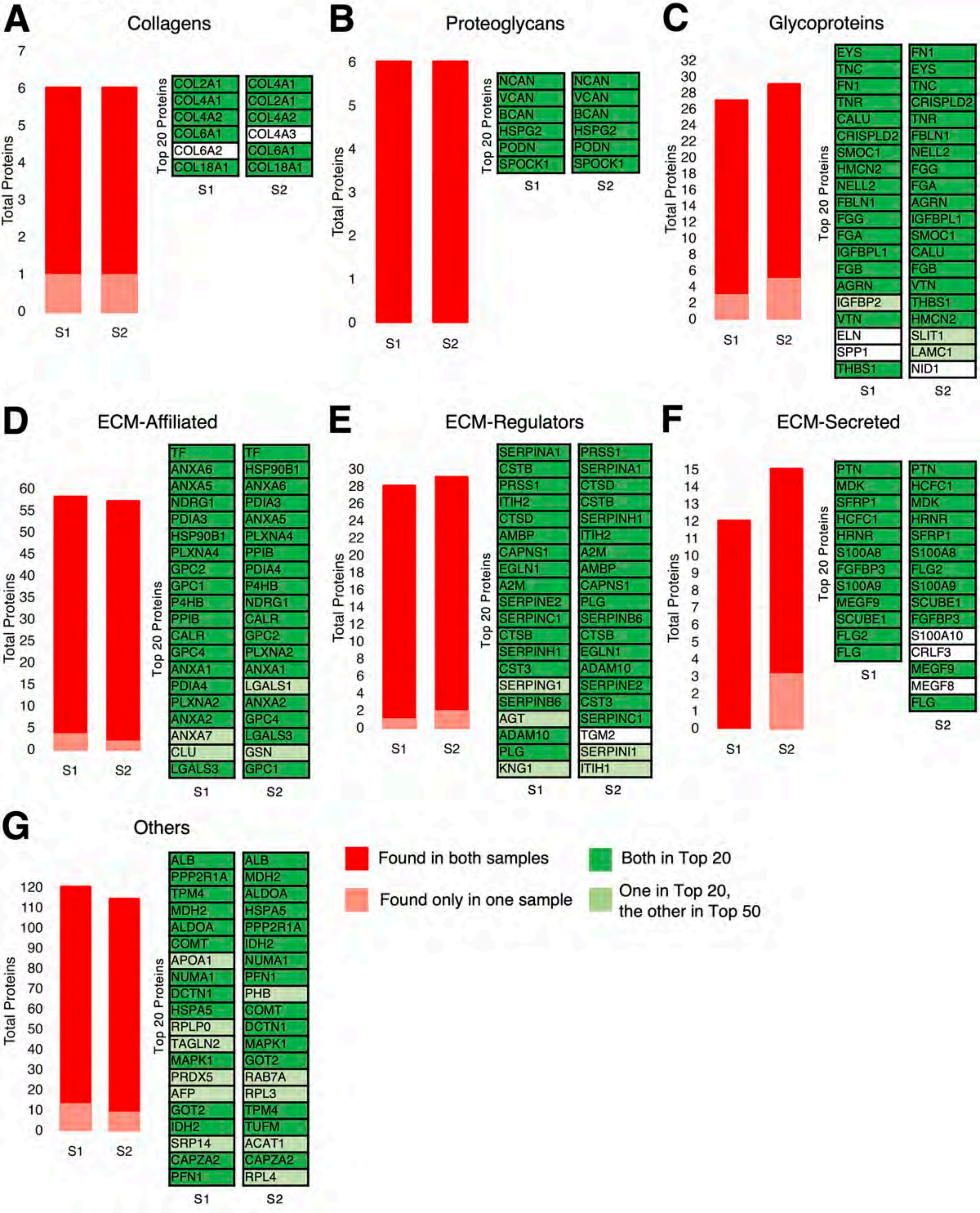
A
Collagens
Total Proteins
Top 20 Proteins
COL2A1
COL4A1
COL4A2
COL6A1
COL6A2
COL18A1
COL4A1
COL2A1
COL4A2
COL4A3
COL6A1
COL18A1
S1
S2
B
Proteoglycans
NCAN
VCAN
BCAN
HSPG2
PODN
SPOCK1
C
Glycoproteins
EYS
TNC
FN1
TNR
CALU
CRISPLD2
SMOC1
HMCN2
NELL2
FBLN1
FGG
FGA
IGFBPL1
FGB
AGRN
IGFBP2
VTN
ELN
SPP1
THBS1
FN1
EYS
TNC
CRISPLD2
TNR
FBLN1
NELL2
FGG
FGA
AGRN
IGFBPL1
SMOC1
CALU
FGB
VTN
THBS1
HMCN2
SLIT1
LAMC1
NID1
D
ECM-Affiliated
TF
ANXA6
ANXA5
NDRG1
PDIA3
HSP90B1
PLXNA4
GPC2
GPC1
P4HB
PPIB
CALR
GPC4
ANXA1
PDIA4
PLXNA2
ANXA2
ANXA7
CLU
LGALS3
TF
HSP90B1
ANXA6
PDIA3
ANXA5
PLXNA4
PPIB
PDIA4
P4HB
NDRG1
CALR
GPC2
PLXNA2
ANXA1
LGALS1
ANXA2
GPC4
LGALS3
GSN
GPC1
E
ECM-Regulators
SERPINA1
CSTB
PRSS1
ITIH2
CTSD
AMBP
CAPNS1
EGLN1
A2M
SERPINE2
SERPINC1
CTSB
SERPINH1
CST3
SERPING1
SERPINB6
AGT
ADAM10
PLG
KNG1
PRSS1
SERPINA1
CTSD
CSTB
SERPINH1
ITIH2
A2M
AMBP
CAPNS1
PLG
SERPINB6
CTSB
EGLN1
ADAM10
SERPINE2
CST3
SERPINC1
TGM2
SERPINI1
ITIH1
F
ECM-Secreted
PTN
MDK
SFRP1
HCFC1
HRNR
S100A8
FGFBP3
S100A9
MEGF9
SCUBE1
FLG2
FLG
PTN
HCFC1
MDK
HRNR
SFRP1
S100A8
FLG2
S100A9
SCUBE1
FGFBP3
S100A10
CRLF3
MEGF9
MEGF8
FLG
G
Others
ALB
PPP2R1A
TPM4
MDH2
ALDOA
COMT
APOA1
NUMA1
DCTN1
HSPA5
RPLP0
TAGLN2
MAPK1
PRDX5
AFP
GOT2
IDH2
SRP14
CAPZA2
PFN1
ALB
MDH2
ALDOA
HSPA5
PPP2R1A
IDH2
NUMA1
PFN1
PHB
COMT
DCTN1
MAPK1
GOT2
RAB7A
RPL3
TPM4
TUFM
ACAT1
CAPZA2
RPL4
Found in both samples
Found only in one sample
Both in Top 20
One in Top 20,
the other in Top 50

Suppl. Fig. 2

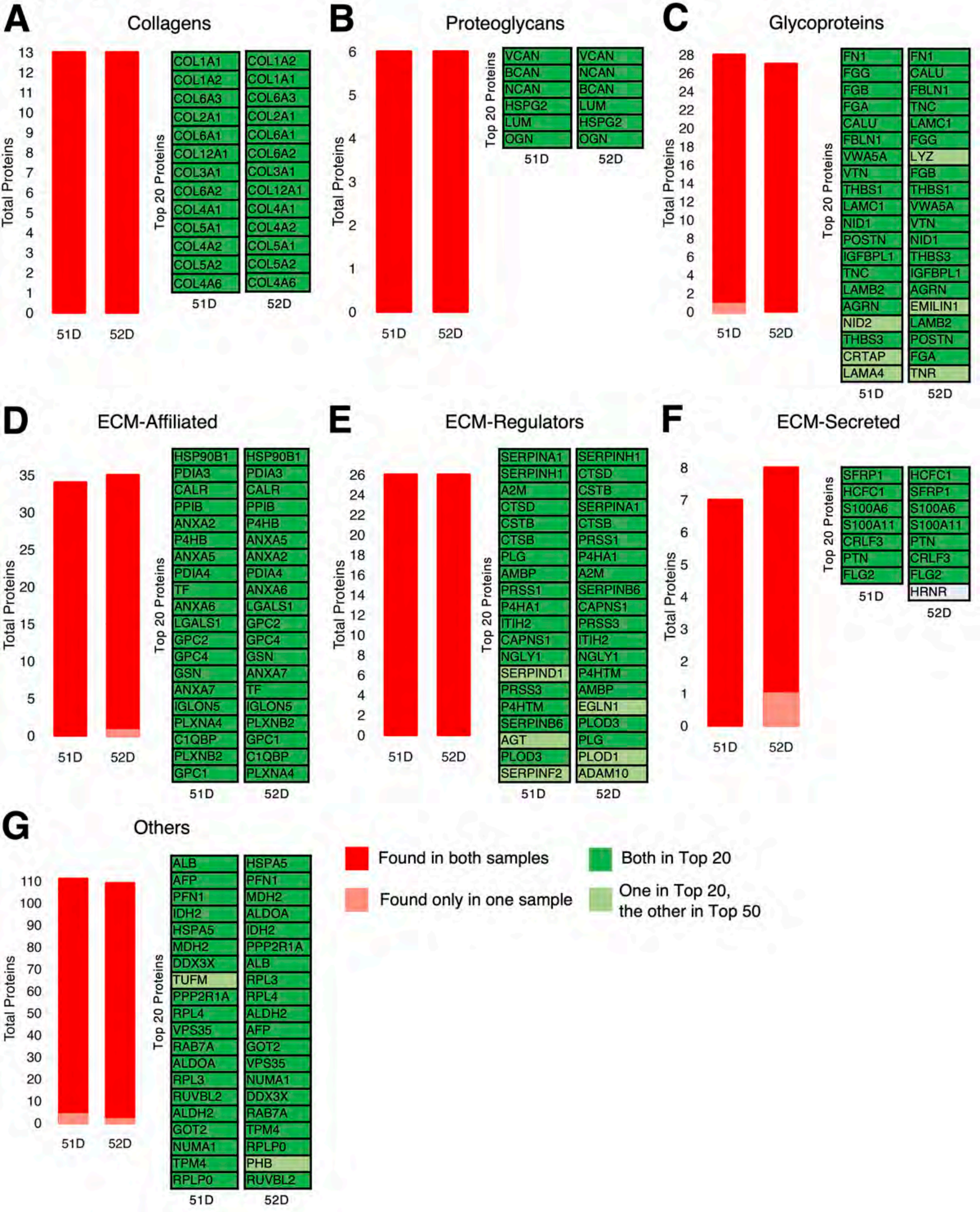

A
Collagens
Total Proteins
13
12
11
10
9
8
7
6
5
4
3
2
1
0
51D
52D
Top 20 Proteins
COL1A1
COL1A2
COL6A3
COL2A1
COL6A1
COL12A1
COL3A1
COL6A2
COL4A1
COL5A1
COL4A2
COL5A2
COL4A6
51D
COL1A2
COL1A1
COL6A3
COL2A1
COL6A1
COL6A2
COL3A1
COL12A1
COL4A1
COL4A2
COL5A1
COL5A2
COL4A6
52D
B
Proteoglycans
Total Proteins
6
5
4
3
2
1
0
51D
52D
Top 20 Proteins
VCAN
BCAN
NCAN
HSPG2
LUM
OGN
51D
VCAN
NCAN
BCAN
LUM
HSPG2
OGN
52D
C
Glycoproteins
Total Proteins
28
26
24
22
20
18
16
14
12
10
8
6
4
2
0
51D
52D
Top 20 Proteins
FN1
FGG
FGB
FGA
CALU
FBLN1
VWA5A
VTN
THBS1
LAMC1
NID1
POSTN
IGFBPL1
TNC
LAMB2
AGRN
NID2
THBS3
CRTAP
LAMA4
51D
FN1
CALU
FBLN1
TNC
LAMC1
FGG
LYZ
FGB
THBS1
VWA5A
VTN
NID1
THBS3
IGFBPL1
AGRN
EMILIN1
LAMB2
POSTN
FGA
TNR
52D
D
ECM-Affiliated
Total Proteins
35
30
25
20
15
10
5
0
51D
52D
Top 20 Proteins
HSP90B1
PDIA3
CALR
PPIB
ANXA2
P4HB
ANXA5
PDIA4
TF
ANXA6
LGALS1
GPC2
GPC4
GSN
ANXA7
IGLON5
PLXNA4
C1QBP
PLXNB2
GPC1
51D
HSP90B1
PDIA3
CALR
PPIB
P4HB
ANXA5
ANXA2
PDIA4
ANXA6
LGALS1
GPC2
GPC4
GSN
ANXA7
TF
IGLON5
PLXNB2
GPC1
C1QBP
PLXNA4
52D
E
ECM-Regulators
Total Proteins
26
24
22
20
18
16
14
12
10
8
6
4
2
0
51D
52D
Top 20 Proteins
SERPINA1
SERPINH1
A2M
CTSD
CSTB
CTSB
PLG
AMBP
PRSS1
P4HA1
ITIH2
CAPNS1
NGLY1
SERPIND1
PRSS3
P4HTM
SERPINB6
AGT
PLOD3
SERPINF2
51D
SERPINH1
CTSD
CSTB
SERPINA1
CTSB
PRSS1
P4HA1
A2M
SERPINB6
CAPNS1
PRSS3
ITIH2
NGLY1
P4HTM
AMBP
EGLN1
PLOD3
PLG
PLOD1
ADAM10
52D
F
ECM-Secreted
Total Proteins
8
7
6
5
4
3
2
1
0
51D
52D
Top 20 Proteins
SFRP1
HCFC1
S100A6
S100A11
CRLF3
PTN
FLG2
51D
HCFC1
SFRP1
S100A6
S100A11
PTN
CRLF3
FLG2
HRNR
52D
G
Others
Total Proteins
110
100
90
80
70
60
50
40
30
20
10
0
51D
52D
Top 20 Proteins
ALB
AFP
PFN1
IDH2
HSPA5
MDH2
DDX3X
TUFM
PPP2R1A
RPL4
VPS35
RAB7A
ALDOA
RPL3
RUVBL2
ALDH2
GOT2
NUMA1
TPM4
RPLP0
51D
HSPA5
PFN1
MDH2
ALDOA
IDH2
PPP2R1A
ALB
RPL3
RPL4
ALDH2
AFP
GOT2
VPS35
NUMA1
DDX3X
RAB7A
TPM4
RPLP0
PHB
RUVBL2
52D
Found in both samples
Found only in one sample
Both in Top 20
One in Top 20, the other in Top 50

Suppl. Fig. 3

### Glycoproteins

| CP | GZ | SP |
|---|---|---|
| EYS | EYS | EYS |
| FN1 | FN1 | FN1 |
| TNR | CALU | TNC |
| CRISPLD2 | TNC | TNR |
| TNC | TNR | NELL2 |
| NELL2 | FBLN1 | FBLN1 |
| FBLN1 | CRISPLD2 | HMCN2 |
| CALU | SMOC1 | CRISPLD2 |
| AGRN | NELL2 | CALU |
| THBS1 | HMCN2 | IGFBP2 |
| HMCN2 | AGRN | SMOC1 |
| IGFBP2 | IGFBPL1 | AGRN |
| SMOC1 | THBS1 | THBS1 |
| VTN | IGFBP2 | SLIT1 |
| FGG | VTN | SPP1 |
| FGA | LAMC1 | IGFBPL1 |
| FGB | ELN | VTN |
| IGFBPL1 | FGG | FGA |
| LAMC1 | FGA | FGG |
| SLIT1 | FGB | ELN |
| SPP1 | LAMB2 | LAMC1 |
| LAMB3 | SLIT1 | VWF |
| CRELD1 | VWF | FGB |
| NID1 | LAMB1 | PCOLCE |
| NID2 | NID1 | IGFBP1 |

### ECM-Affiliated

| CP | GZ | SP |
|---|---|---|
| TF | ANXA6 | TF |
| NDRG1 | PDIA3 | NDRG1 |
| PDIA3 | TF | PDIA3 |
| HSP90B1 | ANXA5 | PLXNA4 |
| ANXA6 | PLXNA4 | ANXA6 |
| PPIB | NDRG1 | HSP90B1 |
| PLXNA4 | PPIB | PPIB |
| P4HB | HSP90B1 | ANXA5 |
| CALR | GPC2 | P4HB |
| ANXA5 | P4HB | PLXNA2 |
| PLXNA2 | ANXA2 | CALR |
| GPC2 | LGALS3 | PLXNA1 |
| PDIA4 | GPC1 | GPC2 |
| PLXNB2 | PLXNA2 | LGALS3 |
| PLXNA1 | ANXA1 | PDIA4 |
| ANXA2 | CALR | GPC1 |
| GPC1 | GPC4 | ANXA2 |
| ANXA1 | PLXNB2 | CLU |
| IGLON5 | PDIA4 | PLXNB2 |
| LGALS3 | ANXA7 | PTGDS |
| ANXA7 | LGALS1 | HPX |
| CLU | CLU | IGLON5 |
| TXNDC5 | IGLON5 | LGALS1 |
| DEFA1 | PLXNA1 | GSN |
| LMAN1 | GSN | ANXA1 |

### ECM-Regulators

| CP | GZ | SP |
|---|---|---|
| SERPINA1 | SERPINA1 | SERPINA1 |
| CTSD | CSTB | CSTB |
| CSTB | PRSS1 | PRSS1 |
| ITIH2 | AMBP | ITIH2 |
| PRSS1 | CTSD | A2M |
| EGLN1 | CAPNS1 | CTSD |
| CTSB | SERPINH1 | EGLN1 |
| A2M | ITIH2 | AMBP |
| SERPINH1 | CTSB | SERPINE2 |
| AMBP | SERPINE2 | CAPNS1 |
| CAPNS1 | EGLN1 | SERPINH1 |
| SERPINE2 | A2M | CTSB |
| CST3 | SERPINC1 | CST3 |
| SERPINI1 | CST3 | SERPINB6 |
| ADAM10 | SERPINB6 | ADAM10 |
| SERPING1 | PLG | SERPINC1 |
| TGM2 | SERPING1 | SERPING1 |
| SERPINA3 | ADAM10 | PLG |
| SERPINC1 | TGM2 | APOE |
| CFH | KNG1 | SERPINI1 |
| PLG | SERPINA3 | KNG1 |
| APOE | AGT | TIMP2 |
| SERPINB6 | SERPINI1 | AGT |
| AGT | CFH | CFH |
| KNG1 | APOE | ADAM23 |

### Others

| CP | GZ | SP |
|---|---|---|
| ALB | ALB | ALB |
| MDH2 | ALDOA | ALDOA |
| PPP2R1A | PPP2R1A | MDH2 |
| HSPA5 | MDH2 | COMT |
| TPM4 | TPM4 | TPM4 |
| GOT2 | NUMA1 | APOA1 |
| ALDOA | PFN1 | PPP2R1A |
| APOA1 | DCTN1 | PFN1 |
| COMT | TAGLN2 | AFP |
| NUMA1 | COMT | IDH2 |
| DCTN1 | MAPK1 | TAGLN2 |
| IDH2 | IDH2 | DCTN1 |
| RPL4 | GOT2 | NUMA1 |
| AFP | HSPA5 | HSPA5 |
| RPLP0 | RUVBL2 | MAPK1 |
| ACAT1 | APOA1 | CAPZA2 |
| PRDX5 | CAPZA2 | GOT2 |
| TUFM | RPLP0 | RUVBL2 |
| RPL3 | PHB | GNA13 |
| CAPZA2 | PRDX5 | RAB7A |
| MAPK1 | CMPK1 | ACAT1 |
| SRP14 | VPS35 | RPL4 |
| PHB | SRP14 | PRDX5 |
| RAB7A | RPL4 | CMPK1 |
| PFN1 | ACAT1 | SRP14 |

### Collagens

| CP | GZ | SP |
|---|---|---|
| COL2A1 | COL4A1 | COL2A1 |
| COL4A1 | COL4A2 | COL1A1 |
| COL4A2 | COL2A1 | COL6A1 |
| COL6A2 | COL4A6 | COL4A2 |
| COL4A6 | COL4A3 | COL1A2 |
| COL4A3 | COL6A1 | COL18A1 |
| COL18A1 | COL6A2 | COL4A1 |
| COL6A1 | COL18A1 | COL6A2 |
| COL1A1 | | |

### Proteoglycans

| CP | GZ | SP |
|---|---|---|
| NCAN | VCAN | NCAN |
| VCAN | NCAN | VCAN |
| HSPG2 | HSPG2 | BCAN |
| BCAN | BCAN | HSPG2 |
| PODN | PODN | PODN |
| SPOCK1 | | SPOCK1 |

### ECM-Secreted

| CP | GZ | SP |
|---|---|---|
| PTN | PTN | PTN |
| MDK | MDK | SFRP1 |
| SFRP1 | SFRP1 | MDK |
| HCFC1 | HCFC1 | HRNR |
| S100A9 | HRNR | HCFC1 |
| S100A8 | SCUBE1 | SCUBE1 |
| HRNR | FGFBP3 | FGFBP3 |
| SCUBE1 | FLG2 | S100A9 |
| FGFBP3 | S100A9 | MEGF9 |
| MEGF9 | S100A8 | S100A8 |
| FLG2 | MEGF9 | FLG2 |
| MEGF8 | S100A10 | MEGF8 |
| CRLF3 | CRLF3 | FLG |
| FLG | FLG | |
| | MEGF8 | |

Present in one layer

Present in two layers

Suppl. Fig. 4

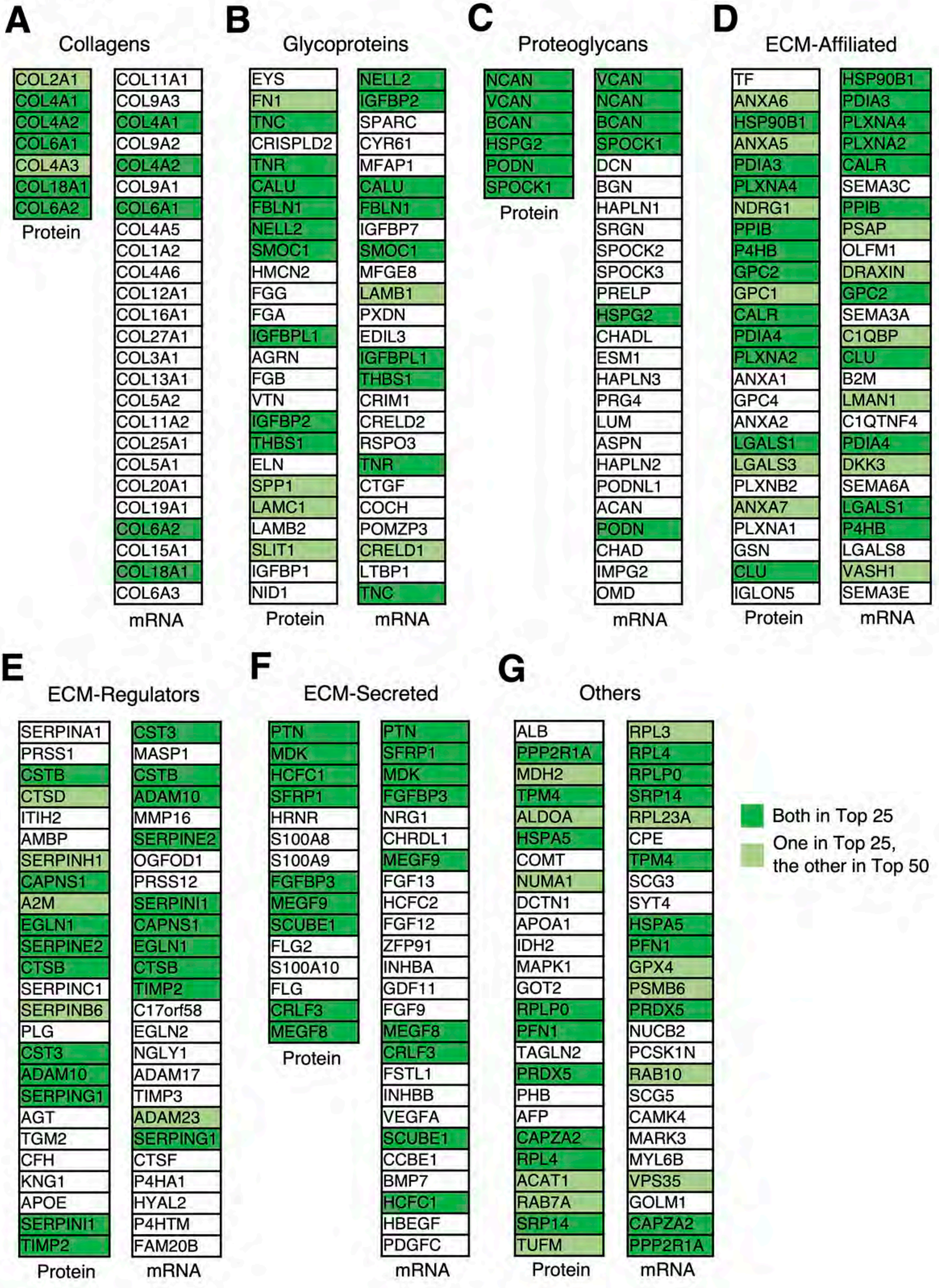
A
Collagens
COL2A1
COL4A1
COL4A2
COL6A1
COL4A3
COL18A1
COL6A2
Protein
COL11A1
COL9A3
COL4A1
COL9A2
COL4A2
COL9A1
COL6A1
COL4A5
COL1A2
COL4A6
COL12A1
COL16A1
COL27A1
COL3A1
COL13A1
COL5A2
COL11A2
COL25A1
COL5A1
COL20A1
COL19A1
COL6A2
COL15A1
COL18A1
COL6A3
mRNA
B
Glycoproteins
EYS
FN1
TNC
CRISPLD2
TNR
CALU
FBLN1
NELL2
SMOC1
HMCN2
FGG
FGA
IGFBPL1
AGRN
FGB
VTN
IGFBP2
THBS1
ELN
SPP1
LAMC1
LAMB2
SLIT1
IGFBP1
NID1
Protein
NELL2
IGFBP2
SPARC
CYR61
MFAP1
CALU
FBLN1
IGFBP7
SMOC1
MFGE8
LAMB1
PXDN
EDIL3
IGFBPL1
THBS1
CRIM1
CRELD2
RSPO3
TNR
CTGF
COCH
POMZP3
CRELD1
LTBP1
TNC
mRNA
C
Proteoglycans
NCAN
VCAN
BCAN
HSPG2
PODN
SPOCK1
Protein
VCAN
NCAN
BCAN
SPOCK1
DCN
BGN
HAPLN1
SRGN
SPOCK2
SPOCK3
PRELP
HSPG2
CHADL
ESM1
HAPLN3
PRG4
LUM
ASPN
HAPLN2
PODNL1
ACAN
PODN
CHAD
IMPG2
OMD
mRNA
D
ECM-Affiliated
TF
ANXA6
HSP90B1
ANXA5
PDIA3
PLXNA4
NDRG1
PPIB
P4HB
GPC2
GPC1
CALR
PDIA4
PLXNA2
ANXA1
GPC4
ANXA2
LGALS1
LGALS3
PLXNB2
ANXA7
PLXNA1
GSN
CLU
IGLON5
Protein
HSP90B1
PDIA3
PLXNA4
PLXNA2
CALR
SEMA3C
PPIB
PSAP
OLFM1
DRAXIN
GPC2
SEMA3A
C1QBP
CLU
B2M
LMAN1
C1QTNF4
PDIA4
DKK3
SEMA6A
LGALS1
P4HB
LGALS8
VASH1
SEMA3E
mRNA
E
ECM-Regulators
SERPINA1
PRSS1
CSTB
CTSD
ITIH2
AMBP
SERPINH1
CAPNS1
A2M
EGLN1
SERPINE2
CTSB
SERPINC1
SERPINB6
PLG
CST3
ADAM10
SERPING1
AGT
TGM2
CFH
KNG1
APOE
SERPINI1
TIMP2
Protein
CST3
MASP1
CSTB
ADAM10
MMP16
SERPINE2
OGFOD1
PRSS12
SERPINI1
CAPNS1
EGLN1
CTSB
TIMP2
C17orf58
EGLN2
NGLY1
ADAM17
TIMP3
ADAM23
SERPING1
CTSF
P4HA1
HYAL2
P4HTM
FAM20B
mRNA
F
ECM-Secreted
PTN
MDK
HCFC1
SFRP1
HRNR
S100A8
S100A9
FGFBP3
MEGF9
SCUBE1
FLG2
S100A10
FLG
CRLF3
MEGF8
Protein
PTN
SFRP1
MDK
FGFBP3
NRG1
CHRDL1
MEGF9
FGF13
HCFC2
FGF12
ZFP91
INHBA
GDF11
FGF9
MEGF8
CRLF3
FSTL1
INHBB
VEGFA
SCUBE1
CCBE1
BMP7
HCFC1
HBEGF
PDGFC
mRNA
G
Others
ALB
PPP2R1A
MDH2
TPM4
ALDOA
HSPA5
COMT
NUMA1
DCTN1
APOA1
IDH2
MAPK1
GOT2
RPLP0
PFN1
TAGLN2
PRDX5
PHB
AFP
CAPZA2
RPL4
ACAT1
RAB7A
SRP14
TUFM
Protein
RPL3
RPL4
RPLP0
SRP14
RPL23A
CPE
TPM4
SCG3
SYT4
HSPA5
PFN1
GPX4
PSMB6
PRDX5
NUCB2
PCSK1N
RAB10
SCG5
CAMK4
MARK3
MYL6B
VPS35
GOLM1
CAPZA2
PPP2R1A
mRNA
Both in Top 25
One in Top 25, the other in Top 50

Suppl. Fig. 5

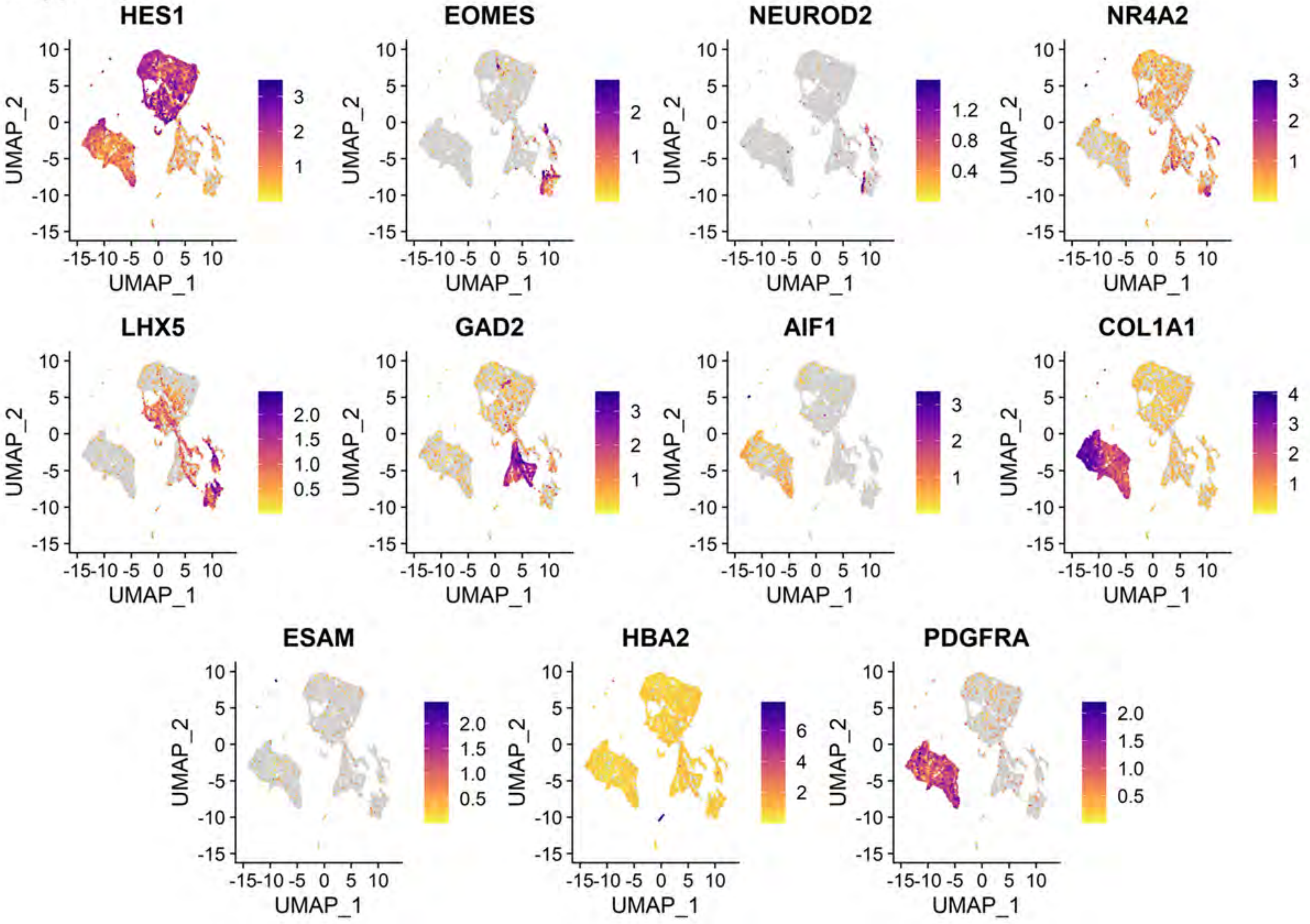
HES1
EOMES
NEUROD2
NR4A2
LHX5
GAD2
AIF1
COL1A1
ESAM
HBA2
PDGFRA
UMAP_1
UMAP_2

Suppl. Fig. 6

**A**

| Collagens | | | |
|---|---|---|---|
| Protein | | RNA | |
| **Gene** | **Abundance** | **Gene** | **Abundance** |
| COL1A2 | 19263979.92 | COL1A2 | 115681 |
| COL1A1 | 15276338.21 | COL3A1 | 73312 |
| COL6A3 | 6416263.333 | COL1A1 | 61048 |
| COL18A1 | 4968346.208 | COL11A1 | 38760 |
| COL5A1 | 4433910.875 | COL5A2 | 35760 |
| COL6A1 | 3703808.625 | COL12A1 | 19777 |
| COL2A1 | 3343561.969 | COL4A2 | 13376 |
| COL12A1 | 2639181.063 | COL6A2 | 13065 |
| COL6A2 | 2615742.5 | COL2A1 | 12142 |
| COL4A2 | 2211137.5 | COL25A1 | 11957 |
| COL11A1 | 1935625.479 | COL26A1 | 10233 |
| | | COL24A1 | 8421 |
| | | COL4A1 | 7754 |
| | | COL6A1 | 6469 |
| | | COL4A5 | 6423 |
| | | COL9A2 | 6387 |
| | | COL5A1 | 6280 |
| | | COL23A1 | 5249 |
| | | COL9A3 | 4793 |
| | | COL18A1 | 4544 |
| | | COL6A3 | 4317 |
| | | COL27A1 | 3728 |
| | | COL21A1 | 3653 |
| | | COL9A1 | 3098 |
| | | COL13A1 | 3096 |

**B**

| Glycoproteins | | | |
|---|---|---|---|
| Protein | | RNA | |
| **Gene** | **Abundance** | **Gene** | **Abundance** |
| FN1 | 66375848.28 | IGFBP2 | 51386 |
| NID1 | 2978040.292 | FBN2 | 34730 |
| TGFBI | 2880291.813 | FN1 | 24949 |
| NID2 | 2757857.208 | FBLN1 | 23663 |
| LAMC1 | 1905357.5 | SPARC | 21384 |
| FGA | 1827712 | HMCN1 | 20922 |
| FGG | 1772995.875 | LTBP1 | 17381 |
| AGRN | 1746605.375 | PCOLCE | 16742 |
| TNC | 1538697.563 | SLIT3 | 12246 |
| FGB | 1305369.792 | LAMA1 | 11654 |
| LAMA1 | 1255636.188 | LAMA2 | 11606 |
| LAMB2 | 1036364.708 | IGFBP5 | 11306 |
| LAMB1 | 898864.4063 | NELL2 | 10734 |
| EMILIN1 | 824739.1146 | IGFBP4 | 10282 |
| EDIL3 | 819409.3125 | CTHRC1 | 9732 |
| LAMA2 | 782590.1563 | LAMA4 | 9728 |
| APOH | 717441.125 | LAMB1 | 9250 |
| IGSF10 | 709282.5 | MFAP4 | 8621 |
| FBLN1 | 649435.875 | PXDN | 8396 |
| MMRN2 | 630172.2813 | SVEP1 | 8246 |
| MFAP1 | 529501.5 | CRIM1 | 7924 |
| AEBP1 | 391253.4063 | SLIT2 | 7536 |
| LAMA4 | 372896.2396 | CRISPLD1 | 7521 |
| THSD4 | 299921.375 | EDIL3 | 7121 |
| VTN | 214016.0938 | EMILIN1 | 6736 |

**C**

| Proteoglycans | | | |
|---|---|---|---|
| Protein | | RNA | |
| Gene | Abundance | Gene | Abundance |
| HSPG2 | 4327001 | LUM | 76526 |
| NYX | 3632028 | VCAN | 41282 |
| VCAN | 3306323.448 | DCN | 29844 |
| LUM | 2533644.667 | OGN | 11430 |
| BGN | 2294178.531 | SPOCK3 | 5657 |
| DCN | 1112050.813 | BGN | 4381 |
| HAPLN3 | 899061.375 | ASPN | 4370 |
| | | SPOCK1 | 3973 |
| | | HSPG2 | 2249 |
| | | HAPLN1 | 2019 |
| | | OMD | 772 |
| | | HAPLN3 | 276 |
| | | PODNL1 | 224 |
| | | PODN | 219 |
| | | ACAN | 160 |
| | | CHADL | 98 |
| | | IMPG2 | 92 |
| | | NCAN | 80 |
| | | PRELP | 75 |
| | | CHAD | 74 |
| | | SRGN | 61 |
| | | SPOCK2 | 54 |
| | | HAPLN4 | 51 |
| | | NYX | 27 |
| | | BCAN | 21 |

**D**

| Regulators | | | |
|---|---|---|---|
| Protein | | RNA | |
| Gene | Abundance | Gene | Abundance |
| SERPINH1 | 39508161.33 | HPSE2 | 59834 |
| SERPINA1 | 7008804.75 | MMP16 | 14569 |
| ITIH2 | 2434454.917 | TIMP3 | 14235 |
| CSTB | 1836127.813 | ADAMTS19 | 13856 |
| TGM2 | 1748372.875 | CST3 | 12933 |
| CTSD | 918467.0938 | CSTB | 11868 |
| CTSC | 776964.6146 | SERPINH1 | 11711 |
| CTSZ | 669489.4375 | ADAMTS6 | 11622 |
| SERPINC1 | 390241.5625 | P4HA1 | 10942 |
| PLOD3 | 343707.0938 | PCSK5 | 9950 |
| P4HA1 | 318829.4063 | ADAMTS9 | 9421 |
| | | ADAMTSL1 | 9262 |
| | | SULF1 | 7917 |
| | | TIMP1 | 7878 |
| | | TIMP2 | 7506 |
| | | MMP2 | 7087 |
| | | CTSK | 6827 |
| | | CTSC | 5906 |
| | | ADAM10 | 5527 |
| | | CTSL | 4970 |
| | | HYAL2 | 4822 |
| | | ADAMTS12 | 4683 |
| | | CPQ | 4604 |
| | | PLOD2 | 4467 |
| | | ADAMTS20 | 4272 |

**E**

| Affiliated | | | |
|---|---|---|---|
| Protein | | RNA | |
| **Gene** | **Abundance** | **Gene** | **Abundance** |
| LGALS1 | 26023453.92 | LGALS1 | 132661 |
| ANXA2 | 20558805.29 | GPC6 | 48117 |
| ANXA6 | 10709966.04 | SEMA3A | 42920 |
| ANXA5 | 8886470.875 | ANXA2 | 32080 |
| ANXA1 | 5582814.75 | COLEC12 | 27757 |
| LGALS3 | 1909588.516 | GPC3 | 22774 |
| LMAN1 | 830733.8438 | PLXDC2 | 18300 |
| COLEC12 | 782255.3125 | SEMA5A | 16893 |
| ANXA4 | 640590.125 | ANXA5 | 14796 |
| ANXA7 | 613515.2188 | ANXA6 | 10356 |
| EMCN | 551678 | UTRN | 9783 |
| PLXND1 | 526861.875 | SDC2 | 9728 |
| CSPG4 | 400810.224 | SEMA3C | 8547 |
| CRTAP | 324766.6563 | LMAN1 | 8271 |
| ANXA11 | 242678.8906 | FREM1 | 7968 |
| | | SEMA6A | 7619 |
| | | PLXNA2 | 7388 |
| | | C1QTNF4 | 7097 |
| | | C1QTNF7 | 4350 |
| | | CRTAP | 3814 |
| | | GPC4 | 3787 |
| | | SEMA3D | 3365 |
| | | ANXA7 | 3308 |
| | | GPC2 | 2616 |
| | | SDC1 | 2583 |

**F**

| Secreted | | | |
|---|---|---|---|
| Protein | | RNA | |
| **Gene** | **Abundance** | **Gene** | **Abundance** |
| S100A11 | 2334709.25 | MDK | 126053 |
| SFRP1 | 1720034.469 | S100A11 | 58111 |
| HCFC1 | 542572.625 | FRZB | 29111 |
| EGF | 380336.4375 | PTN | 26458 |
| FGFBP3 | 242347.7969 | SFRP2 | 22675 |
| HRNR | 55010.23438 | SFRP1 | 20297 |
| | | CXCL14 | 14825 |
| | | CCL2 | 14364 |
| | | IGF2 | 13775 |
| | | FSTL1 | 12492 |
| | | S100A6 | 12315 |
| | | PDGFC | 8158 |
| | | CXCL12 | 7980 |
| | | BMP5 | 7922 |
| | | WNT5B | 7173 |
| | | S100A10 | 6965 |
| | | INHBA | 6840 |
| | | S100A13 | 6649 |
| | | WNT5A | 5745 |
| | | ANGPTL1 | 5636 |
| | | VEGFB | 5010 |
| | | FGF13 | 4499 |
| | | BMP7 | 4373 |
| | | CRLF3 | 4319 |
| | | EGFL6 | 3922 |

G

| Others | | | |
|---|---|---|---|
| Protein | | RNA | |
| Gene | Abundance | Gene | Abundance |
| PPIB | 14447536 | RPL23A | 153593 |
| MIF | 14266026 | RPLP0 | 75580 |
| RPL23A | 9007500.417 | PPIB | 29747 |
| P4HB | 8102621.25 | APOE | 25126 |
| RPLP0 | 7452892.167 | PRDX5 | 22110 |
| IDH2 | 3886921.042 | P4HB | 17867 |
| TTR | 3361564.167 | MYL6B | 17841 |
| PRDX5 | 3113351.667 | PSMB6 | 15604 |
| BDH2 | 2716266.438 | FGFR2 | 14800 |
| PPP2R1A | 2624771.292 | CCDC80 | 14583 |
| CAPZA2 | 2610139.74 | MIF | 14326 |
| PSMB6 | 2429337.125 | CNTN4 | 12692 |
| NUMA1 | 1939696.667 | SVBP | 11004 |
| ARSK | 1723477.75 | DKK2 | 10845 |
| DDX3X | 1651576.396 | MRPL18 | 10834 |
| FGFR2 | 1387300.5 | ACP1 | 9809 |
| RUVBL2 | 1144256.406 | IDH2 | 9632 |
| VPS35 | 1131121.104 | C4orf48 | 9515 |
| HEXB | 1111938.25 | B2M | 9457 |
| HDLBP | 994025.9792 | JAM3 | 9431 |
| F8 | 837306.4375 | DDX3X | 9410 |
| DNM2 | 765607.9844 | NRP2 | 9030 |
| UGGT1 | 742773.4688 | MYDGF | 8865 |
| APOB | 518892.0156 | PTX3 | 8765 |
| HGS | 495180.1563 | CAPZA2 | 8678 |
